# Domoic acid disruption of neurodevelopment and behavior involves altered myelination in the spinal cord

**DOI:** 10.1101/842294

**Authors:** Jennifer M. Panlilio, Neelakanteswar Aluru, Mark E. Hahn

**Affiliations:** Biology Department, Woods Hole Oceanographic Institution, Woods Hole, MA 02543; Massachusetts Institute of Technology (MIT) – Woods Hole Oceanographic Institution (WHOI) Joint Graduate Program in Oceanography and Oceanographic Engineering; Woods Hole Center for Oceans and Human Health; National Institutes of Health / NICHD, Laboratory of Molecular Genetics, Bethesda, MD 20892

**Keywords:** Domoic acid, HAB toxins, developmental toxicity, windows of susceptibility, startle response, myelination

## Abstract

Harmful algal blooms (HABs) produce potent neurotoxins that threaten human health. Early life exposure to low levels of the HAB toxin domoic acid (DomA) produces long-lasting behavioral deficits, but the mechanisms involved are unknown. Using zebrafish, we investigated the developmental window of susceptibility to low doses of DomA and examined cellular and molecular targets. Larvae exposed to DomA at 2 days post-fertilization (dpf), but not at 1 or 4 dpf, showed consistent deficits in startle behavior including reduced responsiveness and altered kinematics. Similarly, myelination in the spinal cord was disorganized after exposure only at 2 dpf. Time-lapse imaging revealed disruption of the initial stages of myelination. DomA down-regulated genes required for maintaining myelin structure and the axonal cytoskeleton. These results identify a developmental window of susceptibility to DomA-induced behavioral deficits involving altered gene expression and disrupted myelin structure, and establish a zebrafish model for investigating the underlying mechanisms.

**GRAPHICAL ABSTRACT:** 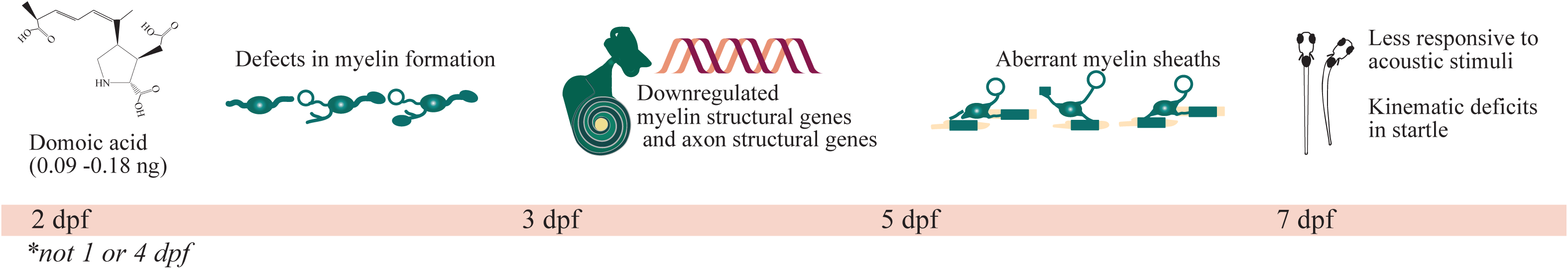

## INTRODUCTION

Domoic acid (DomA) is a potent neurotoxin that is produced by diatoms in the genus *Pseudo-nitzschia.* DomA exerts its toxicity by binding to and activating ionotropic glutamate receptors, particularly the α-amino-3-hydroxy-5-methyl-4-isoxazolepropionic acid (AMPA) and kainate (KA) subtypes.^1^ Human exposure to DomA occurs primarily through the consumption of contaminated seafood. Acute exposure to high levels of DomA leads to amnesic shellfish poisoning, with symptoms ranging from mild gastrointestinal issues to memory loss, seizures, coma, and death.^2–4^ To protect adults from these acute effects, regulatory limits of 20 µg DomA per gram of shellfish tissue have been established.^5,6^ However, seafood with measurable levels of DomA below these regulatory limits is still widely harvested and consumed. This may have important public health consequences, especially for exposures that occur during embryonic and early postnatal development when animals are often more sensitive to neurotoxicants.^7–9^

Research in animal models has demonstrated that developing animals are exposed to DomA and more sensitive than adults to DomA. For example, only one-tenth the dose of DomA is required to induce overt behavioral toxicity in postnatal rats compared to adults.^10–12^ Even within the postnatal period, rats are generally more sensitive at earlier postnatal stages.^12^ Both placental transfer and lactation are potential routes of DomA exposure during development. DomA readily crosses the placenta, making its way into the fetal brain and accumulating in fetal fluids.^13,14^ Amniotic fluid can serve as a reservoir for DomA,^15–17^ suggesting that fetuses could experience prolonged exposure to DomA following a single maternal exposure. DomA can also be transferred to breast milk. DomA has been measured in the milk of sea lions consuming DomA-contaminated prey.^18^ In lactating rats injected with DomA, the toxin is detectable in both the maternal plasma and the milk,^21^ and persists in the milk much longer than it does in the plasma.^19^

A wide range of lasting behavioral deficits can occur following either prenatal or postnatal exposure to DomA. These behavioral effects occur even at doses that do not lead to overt signs of toxicity either in mothers (in the case of prenatal exposures) or in the pups themselves (for postnatal exposures). Rodents exposed prenatally to DomA exhibit aberrant exploratory behaviors,^20–22^ subtle motor coordination deficits,^21^ and in some cases deficits in contextual learning.^21,20^ Rodents exposed postnatally display seizures when exposed to novel environments,^11,23^ and also have aberrant drug-seeking behaviors as assessed by nicotine place preference tests.^24,25^

Together, these studies indicate that developmental exposure to DomA leads to lasting behavioral deficits.^20–22^ However, the cellular and molecular mechanisms underlying these deficits are poorly understood. To elucidate these mechanisms, we used zebrafish as a model. Zebrafish have brain structures and sensory-motor pathways that are homologous to those of humans.^26,27^ Furthermore, the transparency of zebrafish embryos and the availability of transgenic lines allow us to directly observe critical cellular processes during early development.^28–31^ Moreover, larval zebrafish have simple behaviors that are driven by well-characterized neural circuits and comprised of known cell types, allowing us to link behavior to the underlying structural and cellular targets.^32,33^

The goal of this study was to identify the behavioral, structural, and transcriptional changes from low-dose exposures to DomA during critical periods in early development. Using intravenous microinjection, we were able to deliver single doses at specific developmental times that spanned late embryonic (1 day post fertilization, or dpf) to larval stages (4 dpf). We used the well-characterized startle response behavior to identify functional effects from domoic acid toxicity. To assess potential structural changes from exposures, transgenic lines that have fluorescently-labeled myelin sheaths were used to assess changes in myelin structures over time. Finally, transcriptional changes resulting from exposures were identified using RNA sequencing.

## RESULTS

### Developmental exposure to DomA at 2 dpf affects responsiveness to auditory/vibrational stimuli

To elucidate the developmental windows of susceptibility to DomA, we established a zebrafish exposure model involving intravenous injection of DomA into embryos or larvae between 1 and 4 dpf, and then assessed molecular, cellular, and behavioral endpoints at later times (3-7 dpf) (*Materials and Methods; Fig. 1A*).

**Figure 1.**
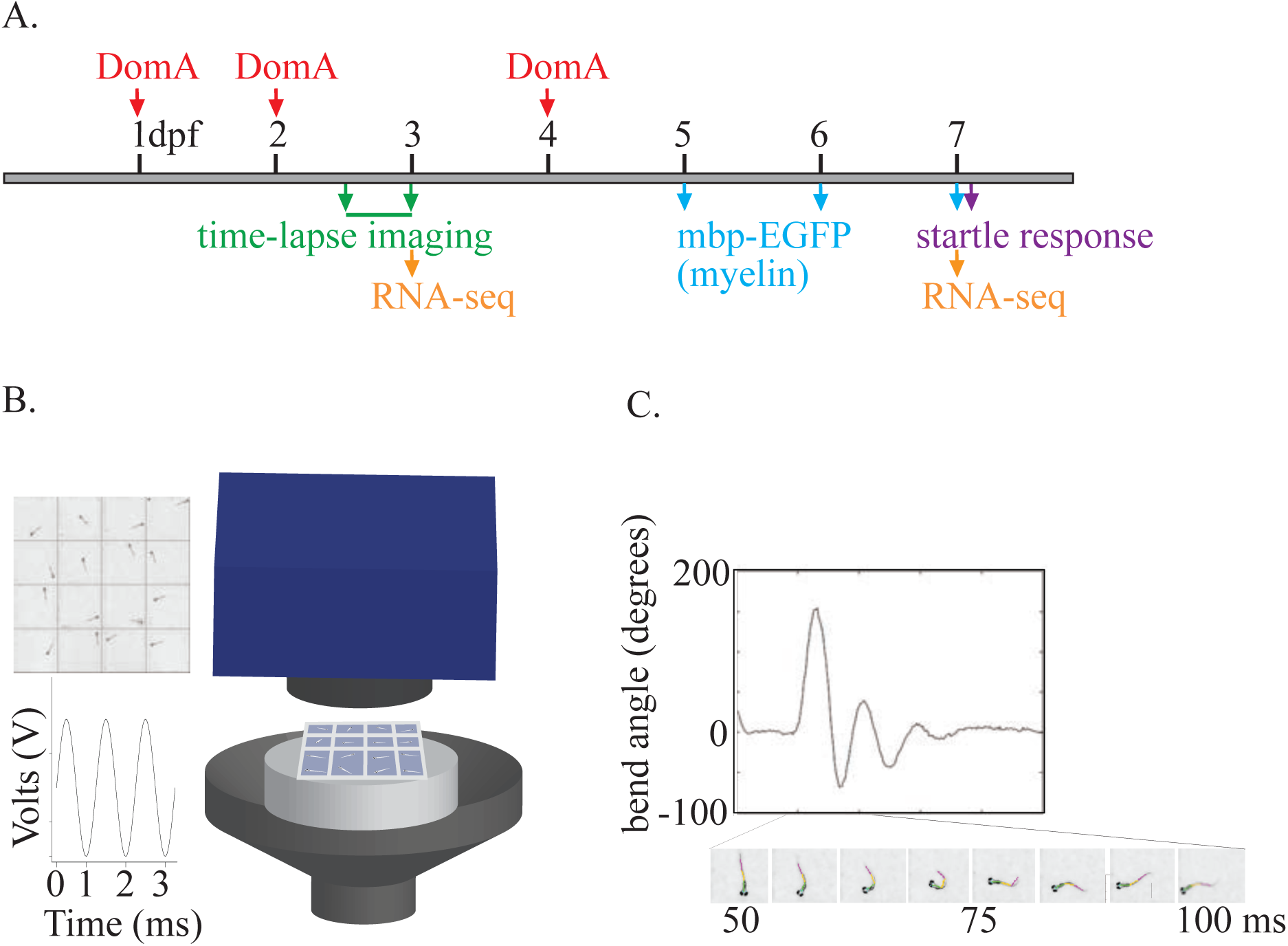
Experimental set-up. **(A)** Exposure paradigm and endpoints assessed over zebrafish development. **(B)** Apparatus used to assess startle responses to A/V stimuli. A speaker with a bonded platform was sent a 3 millisecond, 1000 Hz pulse, which was then delivered to a 16-well plate. A high-speed camera captured startle responses at 1000 frames per second. **(C)** Sample trace of the bend angle over time as a larvae undergoes startle. Bend angle is estimated by measuring the changes in angles between three line segments that outline the larvae.

Injection of DomA at low doses (0.09-0.14 ng) caused transient, acute effects that resolved within one day of exposure and did not lead to appreciable mortality (*Supplemental Results and Discussion; Supplemental Fig.1).*

We assessed the functional impact of developmental DomA exposure by measuring startle response behavior during the larval stage (7 dpf) of development. We first assessed responsiveness—the ability of fish to react to auditory/vibrational (A/V) stimuli—by giving 7 replicate stimuli and calculating the percent response for each fish. Fish exposed to DomA at 2 dpf had reduced responsiveness to A/V stimuli at all doses tested (0.09-0.18 ng) (p <0.001) (Fig. 2). Fish exposed to DomA at 1 dpf had reduced responsiveness when exposed to doses ≥0.13 ng (p ≤0.001), while those exposed to DomA at 4 dpf had reduced responsiveness only when exposed to the highest dose (0.18 ng) tested (p <1e-4). Fish exposed to DomA at 2 dpf were more sensitive than those exposed at 1 or 4 dpf as only fish exposed to DomA at 2 dpf had significantly reduced responsiveness to A/V stimuli at the lowest dose tested (0.09 ng).

**Figure 2.**
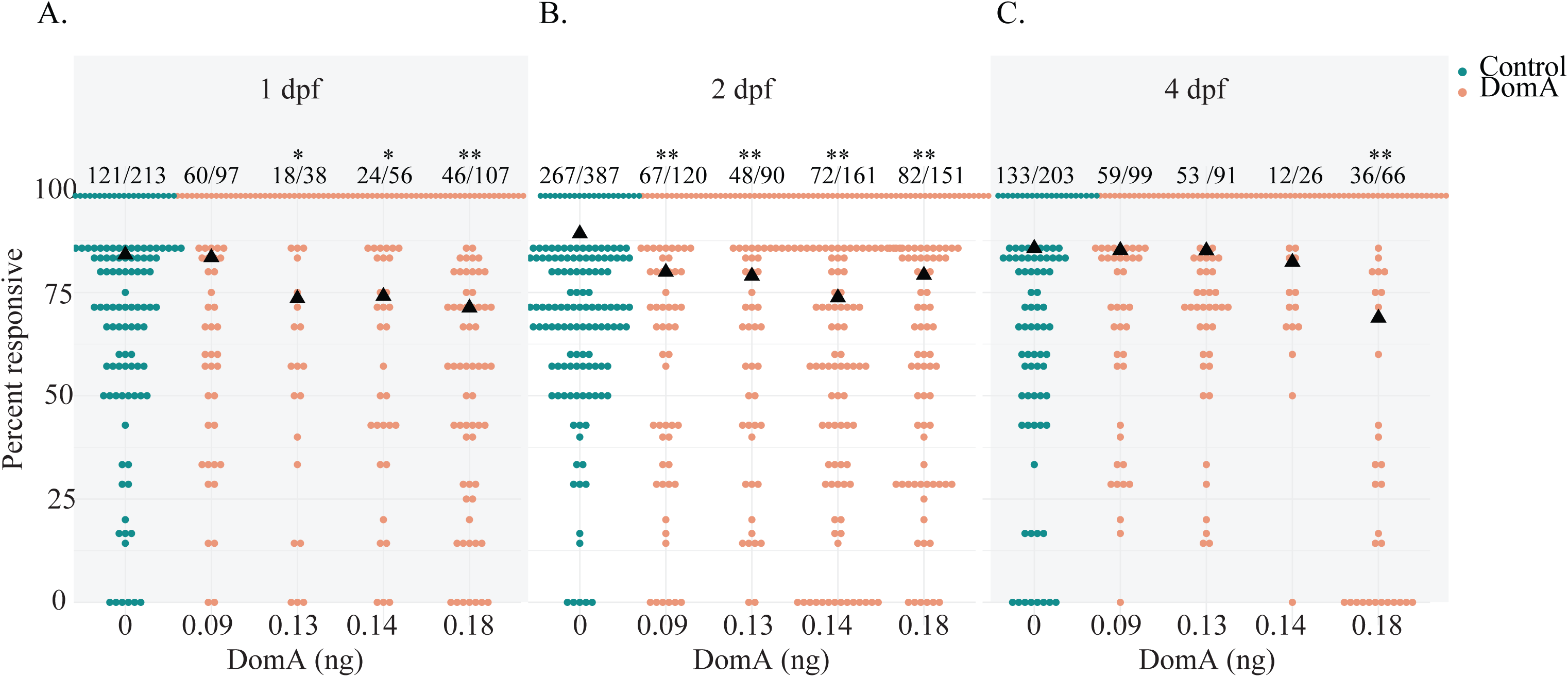
Domoic acid-exposed larvae at 2 dpf are less responsive to auditory/vibrational stimuli. Fish exposed to different doses of DomA at 1 dpf (**A**), 2 dpf (**B**), and 4 dpf (**C**). Ratios listed above represent the number of fish that responded 100% of the time over the total number of fish. Points represent the percent of times an individual fish that responded to replicate stimuli. Black triangles, represent the mean responsiveness of fish for each treatment. Asterisks represent statistical significance between DomA and controls (* p<.05, ** p<.005). Figure supplement: Table 10

### DomA exposure at 2 dpf affects startle response kinematics

During the larval startle response, larvae perform a distinctive ‘c’ bend as the head and body bend together at a high angular velocity (Supplemental Video 1). Kinematics that underlie this ‘c’ bend include bend angle and maximal angular velocity (Mav), which we used to measure DomA-induced changes to startle kinematics. We evaluated kinematics for the two types of startle responses: short latency (SLC) and long latency (LLC) startle responses (Supplemental Fig. 2; *Materials and Methods*).

Exposure to DomA at 2 dpf led to consistent kinematic deficits at all doses tested and in all experimental trials. Fish exposed to DomA at 2 dpf had both reduced bend angles and slower maximal angular velocities relative to vehicle-injected controls; these behavioral deficits were evident with both SLC (Fig. 3) and LLC startle responses (Fig. 4).

**Figure 3.**
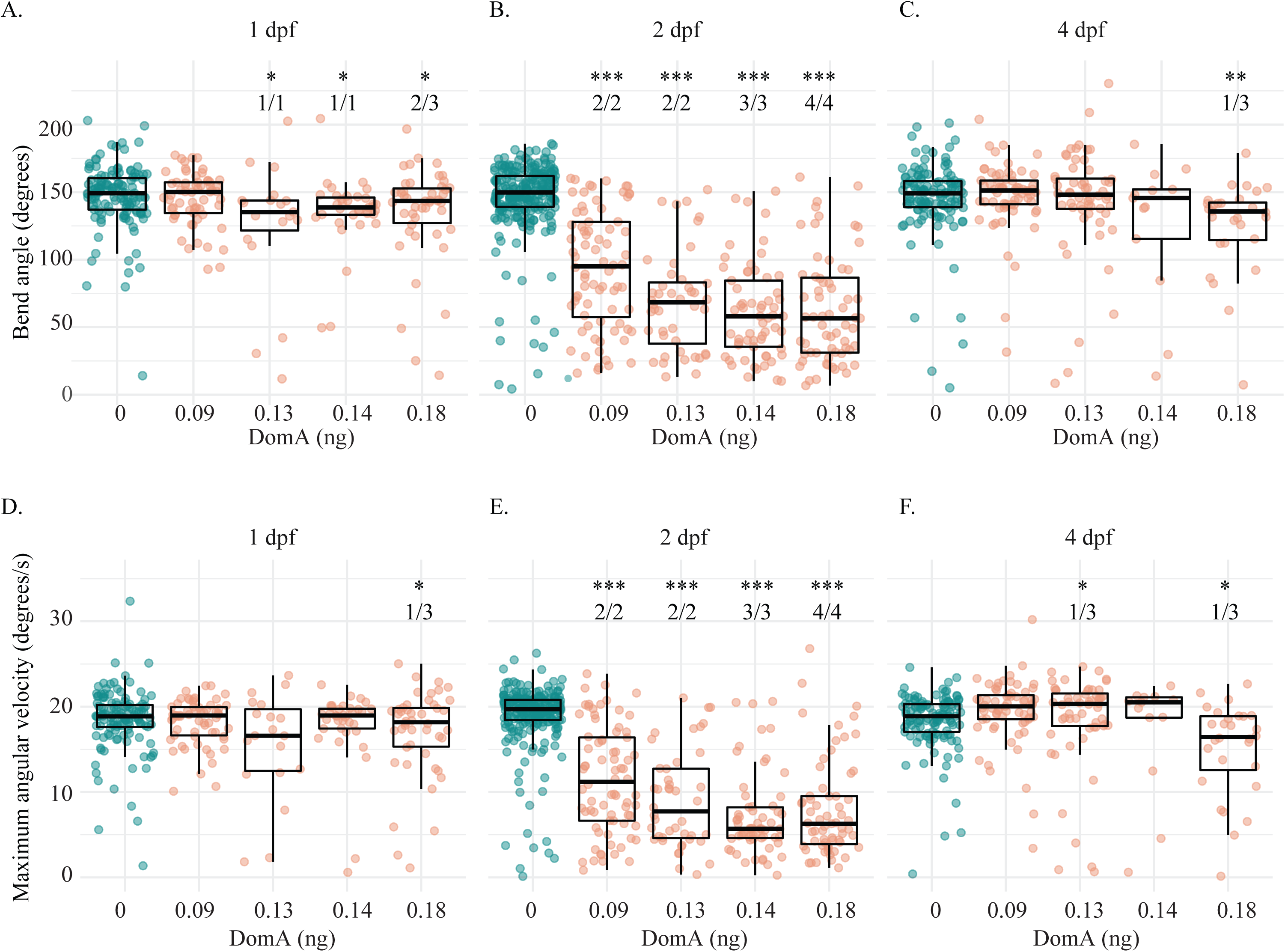
Exposure to domoic acid at 2 dpf (but not 1 or 4 dpf) consistently alters SLC startle response kinematics. Fish were exposed to different doses of DomA at 1 dpf (**A, D**), 2 dpf (**B, E**), and 4 dpf (**C, F**). SLC startle responses were characterized by bend angle (**A-C**) and maximal angular velocity (**D-F**). Each point represents the median of up to 7 responses for an individual fish. Boxplots show the group medians, upper 75% quantiles, and lower 25% quantiles. Asterisks represent statistical significance between DomA and controls (* p < 0.05, ** p <.001, *** p < .0001). The numbers shown above each column represents the number of trials with statistically significant treatment effects / the total number of trials conducted. Figure supplement: Table 11, Table 13 Table 14, Table 16 Table 11, 14 and 16 contains the results from the statistical analysis for 2 dpf, 1 dpf and 4 dpf injected fish. Table 13 includes medians and interquartile ranges for 2 dpf injected fish.

**Figure 4.**
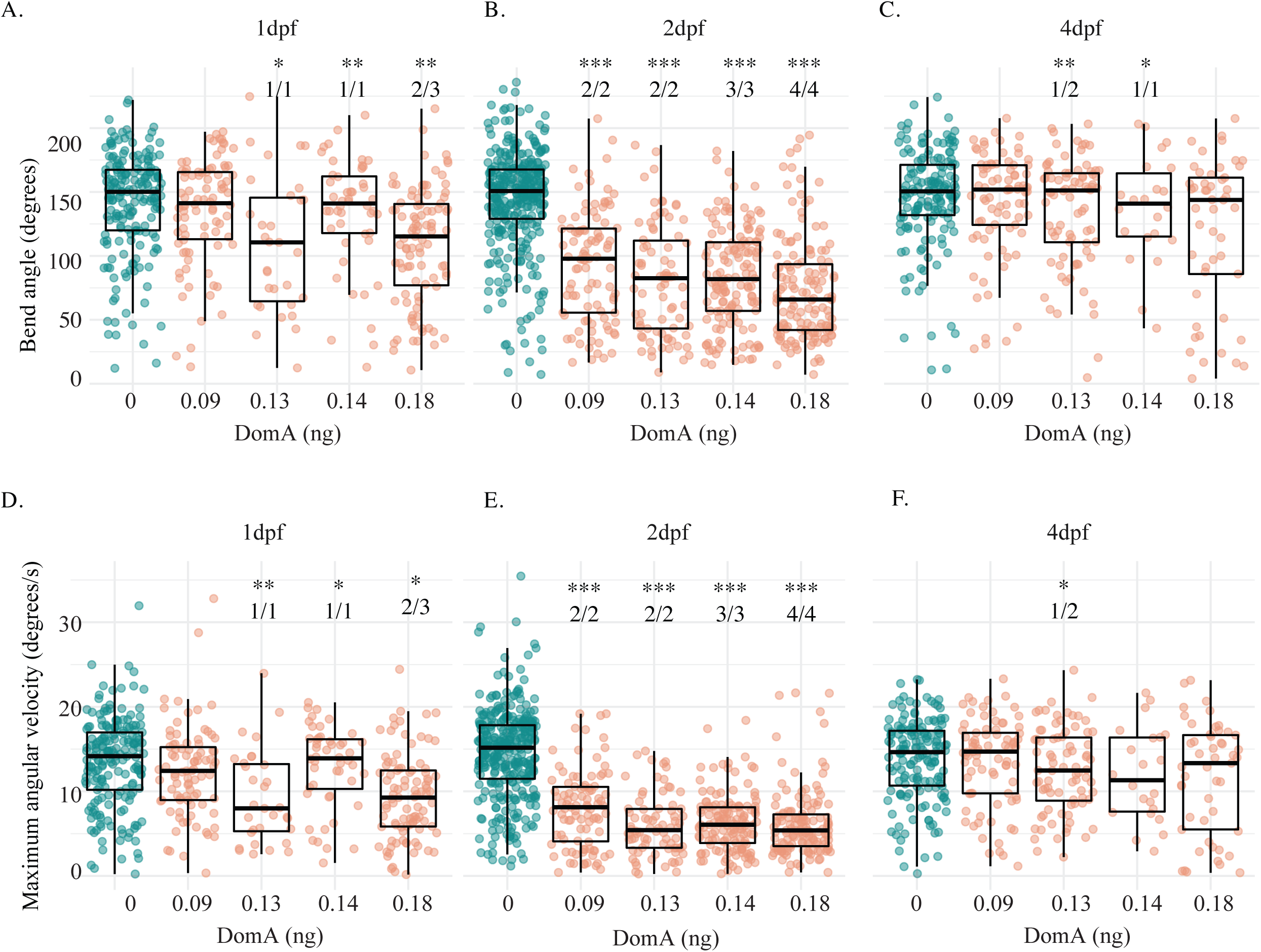
Exposure to domoic acid at 2 dpf (but not 1 or 4 dpf) consistently alters LLC startle response kinematics. Fish were exposed to different doses of DomA at 1 dpf (**A, D**), 2 dpf (**B, E**), and 4 dpf (**C, F**). LLC startle responses were characterized by bend angle (**A-C**) and maximal angular velocity (**D-F**). Each point represents the median of up to 7 responses for an individual fish. Boxplots show the group medians, upper 75% quantiles, and lower 25% quantiles. Asterisks represent statistical significance between DomA and controls (* p < 0.05, ** p <.001, *** p < .0001). The numbers shown above each column represents the number of trials with statistically significant treatment effects / the total number of trials conducted. Figure supplement: Table 12, Table 13, Table 15, Table 17 Table 12, 15, and 17 contains the results from the statistical analysis for 2 dpf, 1 dpf and 4 dpf injected fish. Table 13 includes medians and interquartile ranges for 2 dpf injected fish.

In contrast to exposure at 2 dpf, exposure at 1 and 4 dpf to the lowest dose of DomA tested (0.09 ng) did not lead to any kinematic deficits for either type of startle (SLC or LLC) (Figs. 3, 4). At higher doses (0.13 – 0.18 ng), exposure to DomA at 1 dpf led to kinematic deficits that differed by startle response type. Fish exposed to DomA (≥ 0.13 ng) at 1 dpf had reduced bend angles and slower maximal angular velocities, particularly when they performed the LLC startle responses (Fig. 4). These fish also had significant kinematic deficits when performing the SLC responses, but this was primarily in reductions to bend angle rather than slower maximal angular velocities (Fig. 3). Exposures to DomA at 4 dpf did not result in consistent effects on kinematics. For example, fish exposed to 0.18 ng DomA at 4 dpf had significantly reduced bend angles in only 1 out of 3 trials (Fig. 3). Furthermore, the type of kinematic deficits varied across trials. In 1 of the 3 trials, fish exposed to 0.18 ng DomA had reduced maximum angular velocities and bend angles with SLC startles but not LLC startles. In another trial, fish exposed to DomA at 0.13 ng had deficits in LLC kinematics but not SLC kinematics (Fig. 3 and 4). Thus, while exposures to DomA at all developmental stages tested (1, 2, and 4 dpf) resulted in some kinematic deficits at higher doses, only those at 2 dpf consistently led to kinematic deficits in all trials and across the entire range of doses tested.

To directly compare the effect of both dose and day of exposure on startle kinematics, we performed a nonparametric multivariate factorial analysis on a subset of trials where fish from the same breeding cohort were exposed to DomA at 1, 2, and 4 dpf. We focused on LLC startles because these responses were shown by the previous analysis to be more sensitive to treatment differences. At the lowest dose of DomA (0.09 ng), startle kinematic parameters were significantly influenced by the interaction between treatment and day of exposure (*F*(2, 520)= 21.6, *p* =9.6e-10 for bend angle and − *F*(2, 520)=16.5, *p* =1.1e-7 for Mav) (Supplemental Fig. 3A). Treatment effects from exposure to DomA at 2 dpf were distinct from treatment effects from exposures at 1 or 4 dpf (p < 1e-3). There were no differences in the effects of DomA from exposure at 1 dpf versus 4 dpf, and the kinematics were not significantly different between DomA-exposed fish and their respective controls at these two exposure times. Thus, at the lowest doses of DomA (0.09 ng), exposure at 2 dpf led to distinct kinematic deficits that were not found at 1 or 4 dpf.

With exposure to the intermediate doses of DomA (0.13-0.14 ng), the interaction between treatment and day of exposure remained significant for both bend angle (*F*(2, 474)=23.0, *p*=2.96e-10) and maximal angular velocity (*F*(2, 474)=19.9, *p*=4.84e-9) (Supplemental Fig. 3B). Similar to the results with the lowest dose of DomA, exposure to 0.13 ng DomA at 2 dpf led to significant kinematic deficits relative to exposures at 1 and at 4 dpf (p < 1e-5). Additionally, fish exposed to intermediate doses of DomA at 1 dpf had reduced bend angles and maximum angular velocities, but these deficits were less pronounced compared to those following exposure at 2 dpf (bend angle comparison estimate between 1 dpf – 2 dpf = −140.9 (p = 4.35e-6); maximal angular velocity comparison estimate = −147.92 (p = 1.57e-6)).

### DomA exposure at 2 dpf disrupts myelination in the spinal cord

These startle response deficits could arise from myelin defects. Proper myelination is critical for rapid startle responses, and mutations that disrupt myelin structure cause reduced angular velocities, shallower bend angles, and increased latencies of startle.^34^ To determine whether disrupted myelination underlies the DomA-induced deficits in startle response, we exposed fish with labelled myelin sheaths (*Tg(mbp:EGFP-CAAX)*^35^) to a range of DomA doses and then assessed myelination during the larval stages (Fig. 5A).

**Figure 5.**
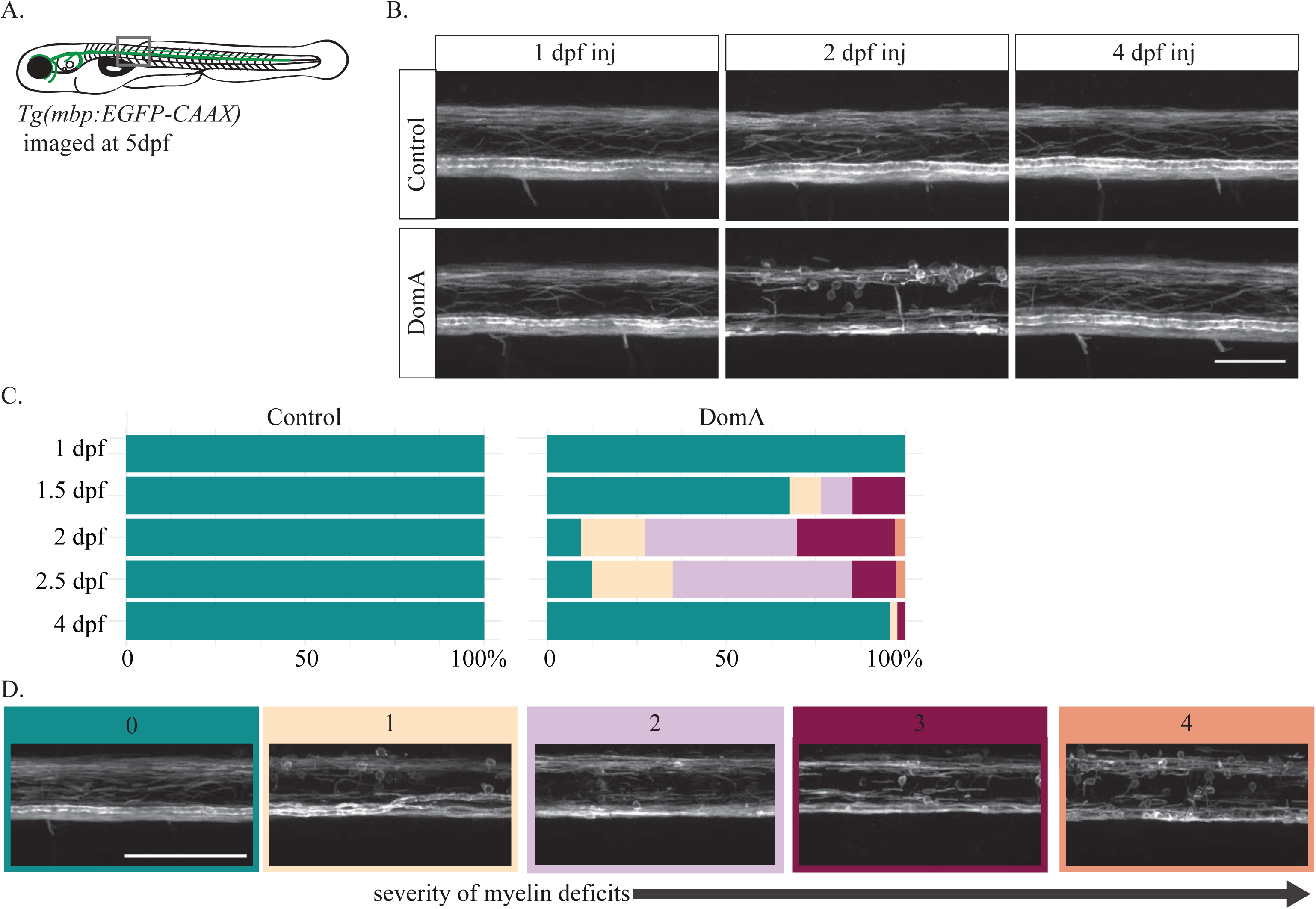
Exposure to domoic acid at 2 dpf (but not 1 or 4 dpf) alters myelin sheaths at 5 dpf. **(A)** *Tg(mbp:EGFP-CAAX)* fish were used to visualize labeled myelin sheaths. **(B)** Fish were exposed to DomA (0.13-0.14 ng) during development (1- 4 dpf), then imaged at 5 dpf using confocal microscopy. Arrows indicate the unusual circular membrane profiles. **(C)** Stacked bar plots show the distribution of the different myelin phenotypes when fish were exposed to DomA at discrete developmental times. Multiple trials were combined to calculate the % distribution per phenotype observed. **(D)** Representative confocal microscopy images of different myelin phenotypes that were observed. Each fish was blindly classified and assigned a category based on severity of the myelin deficit observed. Scale bar = 100 μm. Figure supplement: Table 18 Table 18 includes the number of trials represented along with the associated numbers of fish per trial.

Exposed fish were imaged at 5 dpf using confocal microscopy (Fig. 5B). The severity of myelin defects was scored blindly on the scale of 0-4 (Fig. 5D and Supplemental Fig. 4). Exposure to DomA caused myelin sheath defects, the prevalence and severity of which were influenced by day of exposure (Fig. 5C,D). Fish exposed to DomA at 1 dpf had no visible myelin defects (n= 31). In contrast, 32% of fish exposed at 1.5 dpf had visible myelin defects (n = 11 out of 34). Defects included the overall reduction in labeled myelin, along with the appearance of unusual circular membranes (Fig. 5B). The majority of fish (91%) exposed at 2 dpf showed myelin defects (n =96 out of 106). The prevalence of these defects remained high for fish exposed at 2.5 dpf, with 35 out of 40 (88%) exhibiting a myelin defect. However, these myelin phenotypes were less severe, with 2.5 dpf-exposed larvae having milder myelin sheath defects compared to those exposed to 2 dpf. In comparison, very few fish exposed to DomA at 4 dpf had disrupted myelin sheaths (n= 2 out of 46).

Confocal imaging data suggested that fish exposed at 2 dpf had more severe and more prevalent myelin defects compared to those exposed to DomA at other developmental periods. To confirm this, we performed additional experiments in which fish were exposed to DomA (at various doses and times) and then imaged at 5 dpf using widefield epifluorescence microcopy (Fig. 6). This provided the increased throughput to statistically model the effects of DomA dose and the timing of exposure on the distribution and prevalence of the observed myelin sheath defects.

**Figure 6.**
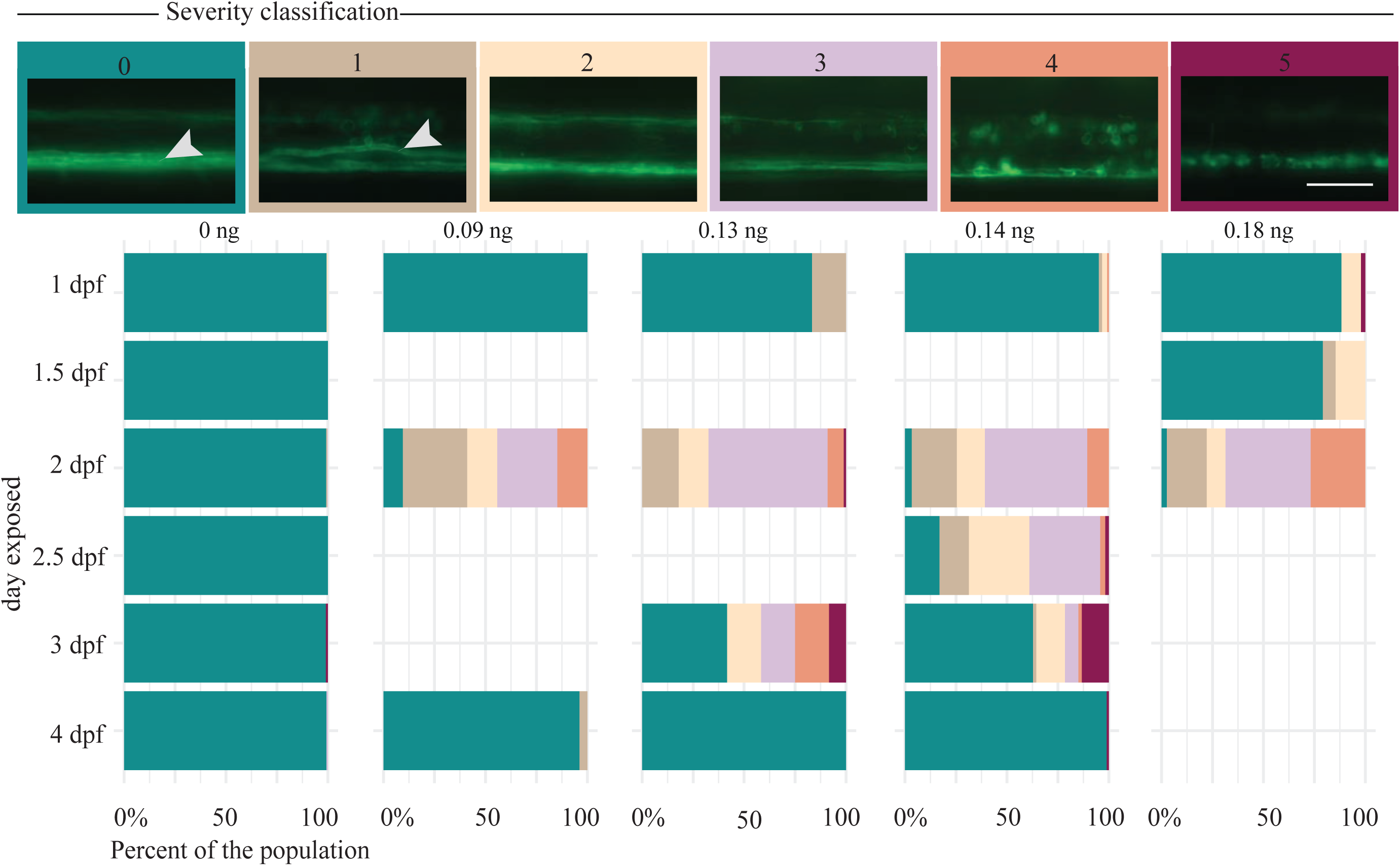
Exposure to domoic acid between 2-2.5 dpf alters myelin sheaths at 5 dpf. **(A)** *Tg(mbp:EGFP-CAAX)* fish were exposed DomA (0.09-0.18 ng) over a range of discrete developmental periods (1-4 dpf), then imaged at 5 dpf using widefield epifluorescence microscopy. Images were blindly classified into 6 categories based on severity of the observed myelin phenotype. Arrows indicate the myelinated Mauthner axon that is required for SLC startle responses. **(B)** Stacked bar plots show the distribution of the different phenotypes. Multiple trials were combined to calculate the % distribution per phenotype observed. Scale bar = 50 μm Figure supplement: Table 19, Table 22, Table 23 Table 19 includes the number of trials represented along with the associated numbers of fish per trial. Table 22 contains the output of the multinomial logistic regression model to assess the role of developmental day of exposure on the distribution of myelin phenotypes. Table 23 contains the output of the multinomial logistic regression model for the influence of dose on the distribution of myelin phenotypes.

To determine whether the day of exposure influenced the appearance and prevalence of myelin defects, we performed a pairwise ANOVA test to compare an initial model, with only DomA dose as the predictor, to an alternative model with both dose and day of exposure as predictors. Incorporating the day of exposure significantly improved its predictive power (p < 1e-16), indicating that timing of DomA exposure influenced myelin deficits.

We then determined whether DomA exposures that occurred during particular periods in development led to a higher prevalence of specific myelin defects. We found that the odds of fish exhibiting phenotypes from category 1-4 were higher when exposures occurred at 2, 2.5, and 3 dpf relative to exposures that occurred at 1 dpf (Supplemental Table 22, p < 1e-7 for 2 dpf exposed). Of these time periods, exposures at 2 dpf had the highest odds of having fish with these myelin defects.

To determine whether these myelin phenotypes observed at 5 dpf persist, fish were also imaged at 6 and 7 dpf (Fig. 7). Similar to imaging at 5 dpf, fish exposed to DomA at 2 dpf and then imaged at 6 or 7 dpf had a significantly higher incidence of myelin defects compared to control fish. Furthermore, the higher the dose of DomA (delivered at 2 dpf), the more likely it was for the fish to exhibit all of the myelin phenotypes observed (Fig. 7A and 7B). These results indicate that DomA exposure, particularly at 2 dpf, leads to myelin defects that persist for at least seven days after exposure.

**Figure 7.**
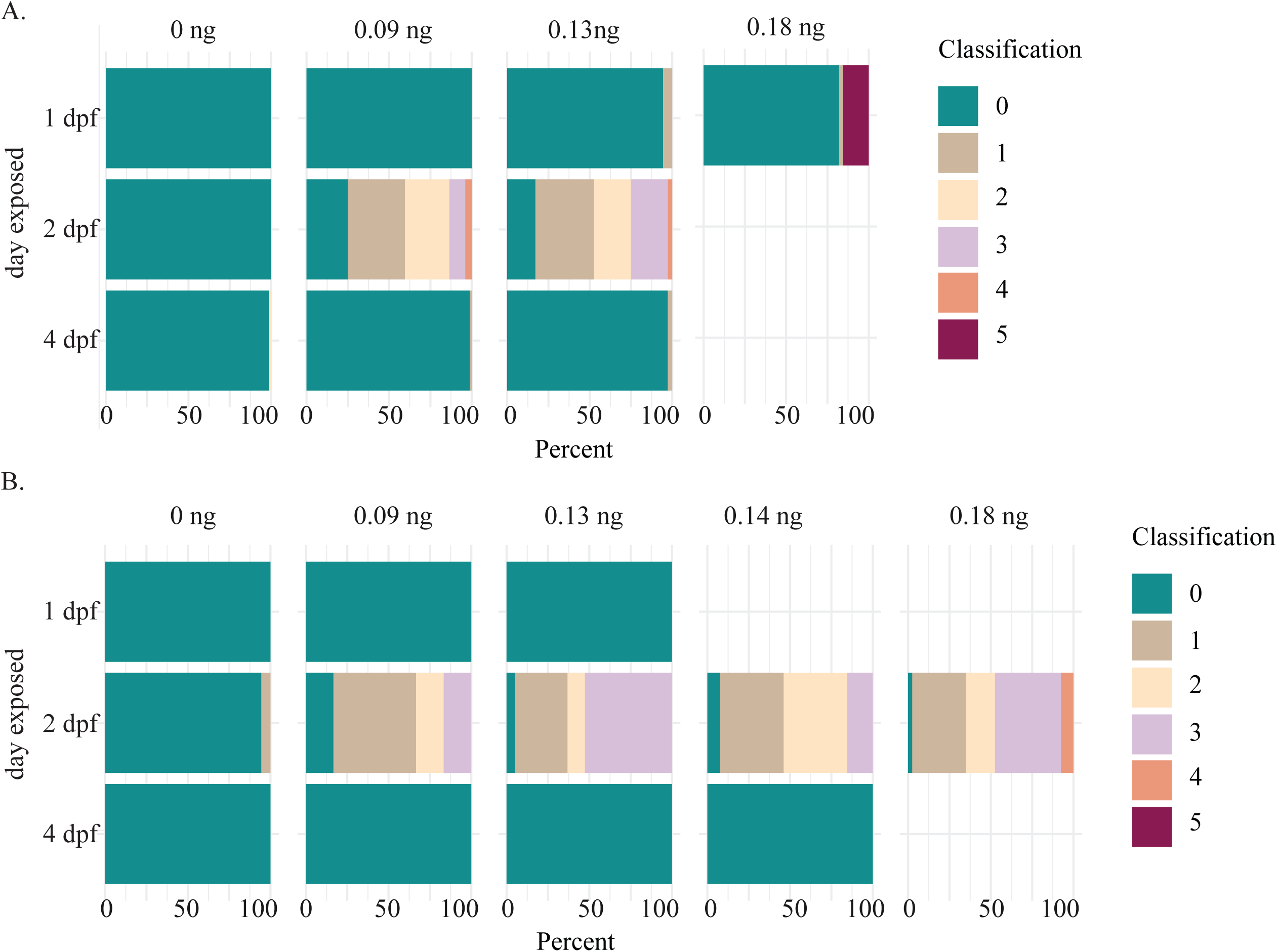
Myelin sheath labeling defects persist until at least 7 dpf. *Tg(mbp:EGFP-CAAX)* fish were exposed to DomA over discrete developmental periods (1, 2 and 4 dpf), then imaged at 6 dpf (**A**) and 7 dpf (**B**) using widefield epifluorescence microscopy. Stacked bar plots show the distribution of the different phenotypes per each dose. Multiple trials were combined to calculate the % distribution per phenotype observed. Figure supplement: Table 20, Table 21, Table 23 Table 20 and 21 contains the number of trials and associated numbers of fish per trial for 6 dpf (Figure 7A) and 7 dpf injected fish (Figure 7B). Table 23 contains the output of the multinomial logistic regression model for the influence of dose on the distribution of myelin phenotypes.

### Time-lapse imaging shows that domoic acid perturbs the initial stages of myelin sheath formation

We observed very few myelination defects or behavioral phenotypes in larvae exposed to DomA at 4 dpf, a time point after the onset of myelination. This suggests that DomA does not affect established sheaths, but rather may perturb the formation of nascent myelin. DomA-exposed fish have perturbed myelin sheaths by 3 dpf (the earliest development period at which myelin sheaths are established) (Fig. 8A). To directly visualize the initial stages of myelin sheath formation, we performed time-lapse imaging in double transgenic fish (*Tg:sox10:RFP*; *Tg:nkx2.2a:mEGFP*), in which cells of the oligodendrocyte lineage—the cells responsible for myelination in the central nervous system—are labeled. Imaging the axon wrapping and nascent myelin sheath formation from 2.5-3 dpf confirmed that oligodendrocytes in DomA-exposed larvae were unable to form elongated sheaths, but rather formed unusual circular membranes (n=5 for controls, n= 6 for DomA exposed larvae) (Fig. 8B, Supplemental Video 2, 3).

**Figure 8.**
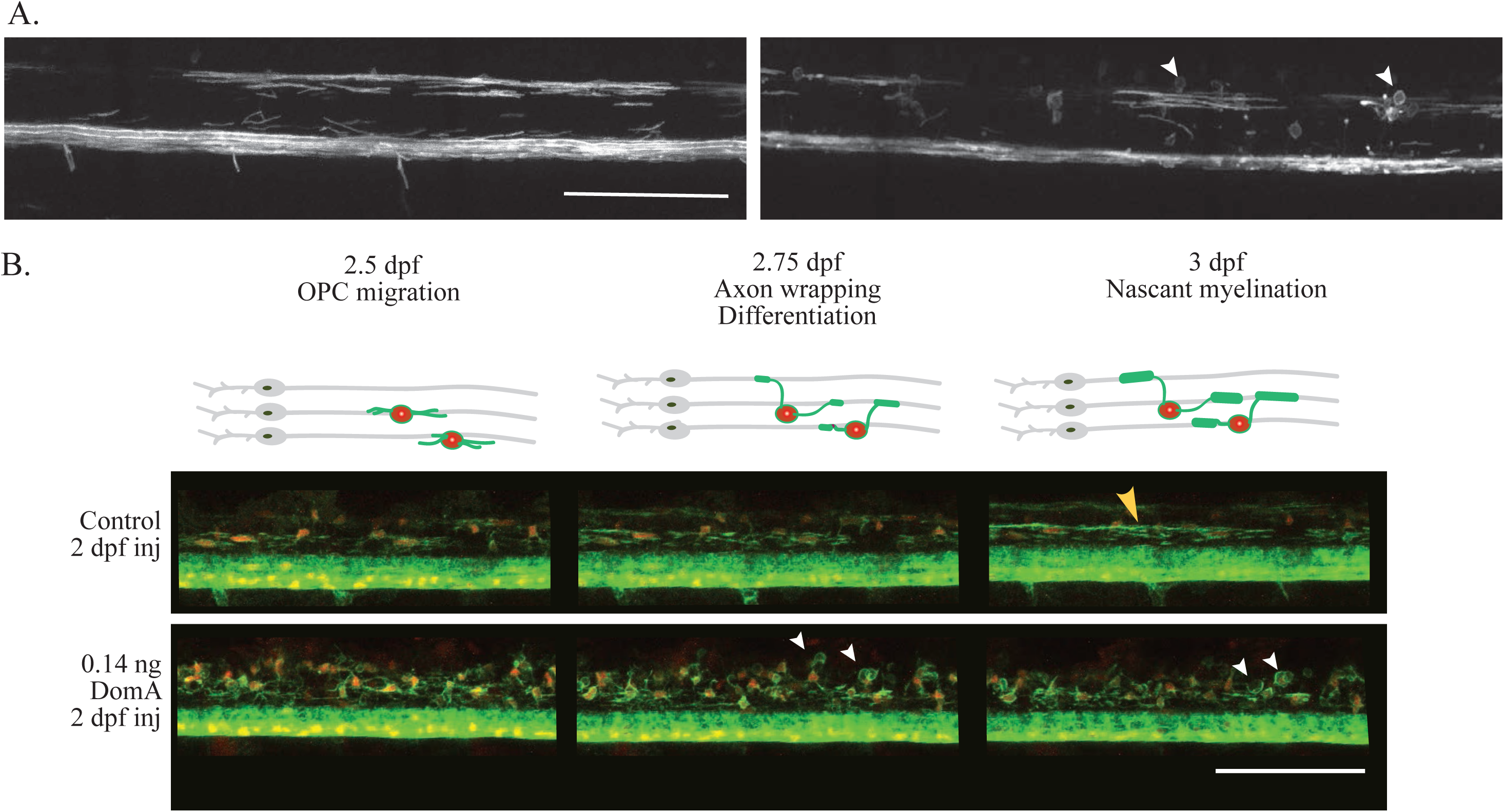
Domoic acid perturbs the initial formation of myelin sheaths. **(A)** *Tg(mbp:EGFP-CAAX)* fish were used to visualize labeled myelin sheaths. Larvae exposed to domoic acid had fewer labeled myelin sheaths compared to controls at the earliest time point myelin sheaths are detected (3 dpf). Furthermore, DomA-exposed larvae also had aberrant circular protrusions by 3 dpf (white arrows) (control, n=5 and DomA, n=10). **(B)** Stills from time-lapse imaging of *Tg(nkx2.2:mEGFP)* x *Tg(sox10:mRFP)* from 2.5- 3 dpf. Diagrams above the images show the key developmental processes in the oligodendrocyte lineage during this time range (control, n=6 and DomA, n=5). Yellow arrow denotes an elongated myelin sheath, white arrows denote unusual circular myelin membranes. Scale bar = 100 μm Figure supplement: Stills (Fig. 8B) were from a time-lapse of control (Supp. video 2) and DomA exposed (Supp. video 3) *Tg(nkx2.2:mEGFP)* x *Tg(sox10:mRFP)* transgenic fish that were imaged from 2.5- 3 dpf.

### Domoic acid exposure alters expression of genes involved in axonal growth and myelination

To identify the gene expression changes that accompany the myelination and startle deficits, whole-embryo RNAseq was performed on embryos exposed to 0.14 ng DomA at 2 dpf and then sampled at 3 and 7 dpf (Fig. 9A).

**Figure 9.**
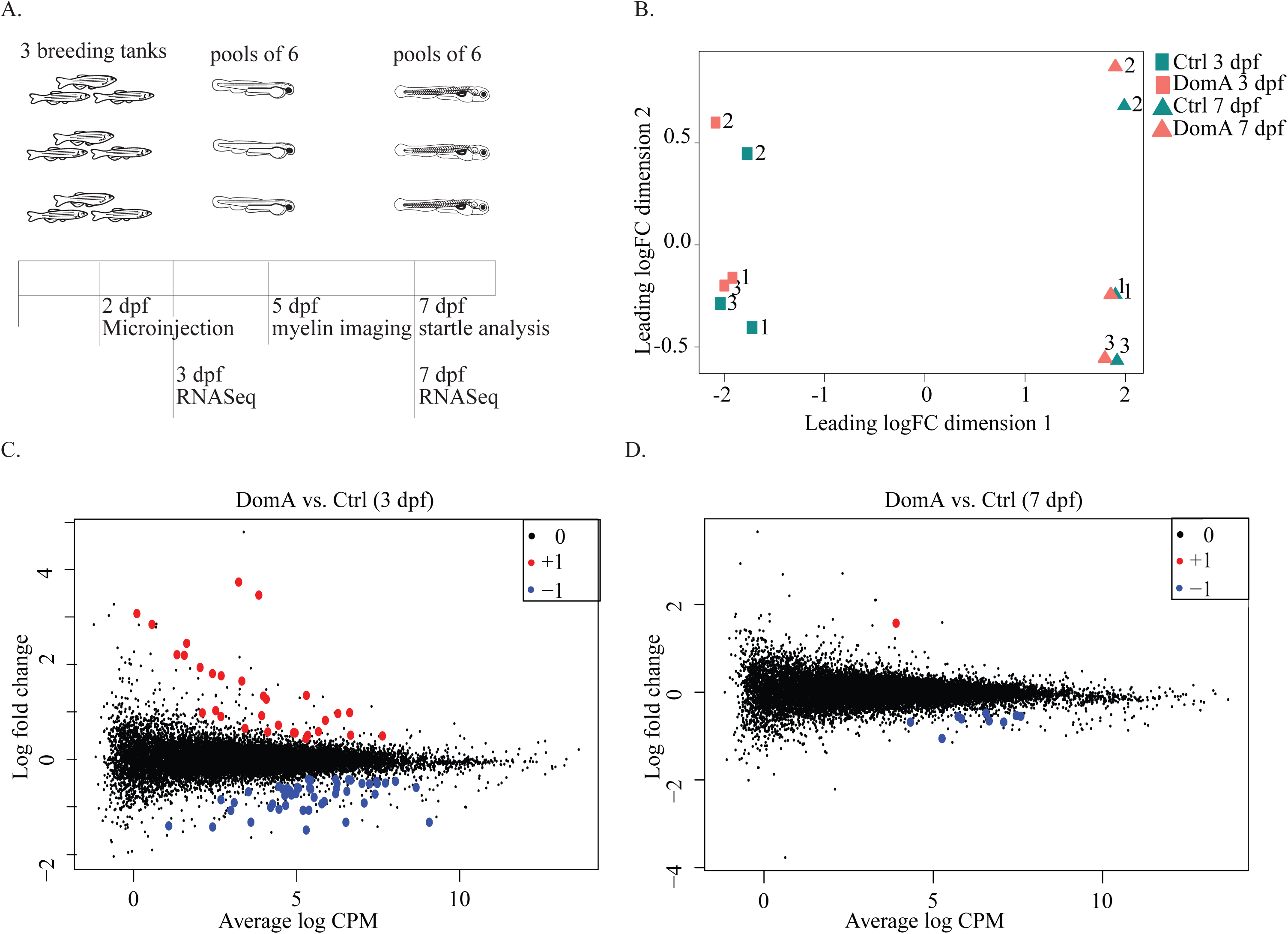
Transcriptional changes associated with domoic acid exposure at 2 dpf. **(A)** Experimental design. Tanks of 3 adult fish of (2 females, 1 male) *Tg(mbp:EGFP-CAAX)* background were bred and exposed to DomA or vehicle at 2 dpf. Pools of 6 embryos within a given treatment from each tank were then sampled at 3 dpf and 7 dpf for RNAsequencing. For functional analyses, myelin sheath labeling was assessed at 5 dpf and startle response was assessed at 7 dpf prior to RNAsequencing. **(B)** MDS plot shows clustering of samples based on overall differences in expression profiles. **(C-D)** Mean-difference (MD) plots compare the log fold changes of genes in DomA exposed versus control fish at the 3 and 7 dpf sampling times. Figure supplement: Table 24, 25, 26, 27

RNA sequencing yielded an average of 21 million raw reads per sample. Of these, 77.6% were uniquely mapped to the zebrafish genome. A multidimensional scaling (MDS) plot revealed clustering by both developmental stage (3 dpf vs. 7 dpf) and breeding clutch (3 breeding trios) (Fig. 9B). This indicates that the differences in gene expression were driven primarily by developmental stage and breeding clutch. However, a number of genes were identified as being differentially expressed in response to DomA.

Statistical analysis revealed differential expression of 82 genes at 3 dpf (28 hours post exposure), and 10 genes at 7 dpf in DomA-exposed fish versus controls (Fig. 9 C, D). Among the 82 genes differentially expressed at 3 dpf, 51 genes were down-regulated and 31 were up-regulated in DomA-exposed larvae as compared to controls.

Pathway analysis of the differentially expressed genes (DEGs; DomA vs. control) indicated an overrepresentation of the GO biological process terms protein depolarization and microtubule depolarization. The genes represented under these GO terms include genes in the stathmin family. Two out of three stathmin genes were up-regulated, and one was down-regulated in DomA-exposed fish.

Significant human phenology phenotypes associated with the down-regulated genes included peripheral axonal degeneration, segmental peripheral demyelination/remyelination, and myelin outfoldings. Several genes required for the maintenance of axonal and myelin structure (*neflb, nefmb, nefma, nefla, mpba, mpz*) were downregulated in DomA-exposed fish relative to controls, and were overrepresented in the human phenology phenotypes (Fig.10). There were no human phenology phenotypes associated with up-regulated genes.

At 7 dpf, there were only ten DEGs, with 9 down-regulated and 1 up-regulated in DomA-exposed fish relative to the controls (Fig. 9D). Comparison of DEGs from 3 and 7 dpf revealed 4 out of the 10 genes to be common to both the time points. Among these, 3 were down-regulated and 1 was up-regulated, with only 2 being annotated. Two of the three shared down-regulated genes were neurofilament genes required for maintaining axonal integrity (*nefmb* and *neflb*).

## DISCUSSION

It is well known that early development is a period of enhanced sensitivity to effects of DomA exposure, and that low-doses of DomA can lead to persistent behavioral deficits^10–12,20–22,24,25^. However, the mechanisms that underlie these changes are largely unknown. This study identified the period around 2 dpf as a window of susceptibility to DomA neurodevelopmental toxicity and then characterized the resulting molecular, structural, and behavioral consequences of exposures during this period. Exposure to DomA during this window led to changes in gene expression, disruption of myelin sheath formation in the spinal cord, and aberrant startle behavior.

### A novel exposure method uncovers a window of susceptibility to low doses of DomA

This study established zebrafish as a model for investigating the mechanisms of toxicity from low-dose exposures to DomA during development. Previous developmental DomA exposure studies in zebrafish were done by injecting DomA into the yolk during the early embryonic stages (512-1000 cell stage).^36,37^ However, the DomA doses that led to behavioral phenotypes were also those that resulted in high mortality rates and lasting neurotoxic symptoms. To build on this work, we used a novel exposure method in which DomA was delivered intravenously at different periods in development – from the embryonic to the larval stages. Using this method, we were able to find a window of susceptibility for low doses of DomA (nominal doses 3-to 260-fold lower than those used previously) at which structural and behavioral effects occurred with no appreciable mortality and minimal gross morphological defects. In particular, the period around 2 dpf was identified as the window of susceptibility for nominal doses of DomA that ranged from 0.09-0.14 ng per embryo.

### Startle response deficits are dependent on dose and timing of exposure

Startle response behavior was used as a functional read out of developmental neurotoxicity. Fish exposed to DomA at 2 dpf (but not 1 and 4 dpf) had aberrant startle behavior at all doses tested (0.09-0.18 ng). In particular, fish exposed to DomA at 2 dpf had reduced responsiveness, increased latency, slower maximal angular velocities, and lower bend angles relative to controls (Figs. 2-4). This suggests that there is a window of susceptibility to low-dose (< 0.18 ng) DomA exposure at around 2 dpf that leads to a functional change in behavior.

### Exposure to DomA at 2 dpf disrupts myelin formation

Similar to the behavioral results, only fish exposed at 2 dpf (but not 1 or 4 dpf) showed consistent defects in myelination within the spinal cord (Figs. 5B,C, 6B). DomA-exposed larvae had an overall reduction in labeled myelin, along with the appearance of unusual circular membranes (Figs. 5B, 6B). These deficits were visible as early as 3 dpf, when nascent myelin sheaths are present (Fig. 8), and persisted until at least 7 dpf, indicating that initial formation of myelin is perturbed and does not recover within 4 days post-exposure (Fig. 7).

### The window of susceptibility to DomA corresponds to the critical period for oligodendrocyte development

It is possible that 2 dpf is the window of susceptibility because DomA perturbs specific developmental processes that occur within this time period. While most of the early neurons have already differentiated by 2 dpf, the oligodendrocyte lineage – the lineage that myelinates axons in the central nervous system – is just beginning to migrate and differentiate during this period.^38,39^ DomA exposure at 2 dpf may perturb critical processes in oligodendrocyte development, leading to the observed disrupted myelination.

Both myelinating oligodendrocytes and their precursors express functional ionotropic glutamate receptors, making them potential cellular targets for DomA.^40,41^ Previous studies have shown that kainate, a structural analog of DomA, causes cell death in oligodendrocyte primary cell cultures, at concentrations comparable to those affecting neurons.^42–45^ Binding to and activating AMPA receptors inhibits the proliferation and differentiation of oligodendrocyte precursor cells into mature oligodendrocytes *in vitro*.^46,47^ Mature oligodendrocytes have also been shown to undergo demyelination after chronic direct infusion of kainate on the optic nerves.^48^ All of this suggests that DomA may alter oligodendrocyte development, and that exposure to DomA at 2 dpf may disrupt critical processes important for OPC proliferation, differentiation, or myelin sheath formation.

Only one previous study has assessed myelin following developmental exposure to DomA. Eleven week-old juvenile mice exposed *in utero* during gestational days 11.5 and 14.5, but not 17.5, had a reduced staining for the myelin-associated glycoprotein (MAG) in their cerebral cortices.^20^ This suggests that there may be periods in early development that are more sensitive to exposure to DomA, leading to these myelination deficits. Indeed, it is possible that sensitivity at the early periods is due to disruptions in oligodendrocyte development, thereby altering their ability to form myelin sheaths during the postnatal period.^49,50^ Our findings extend this work by identifying altered myelination in the spinal cord and revealing that DomA does not disrupt already established myelin sheaths but rather perturbs the initial formation of the sheaths during a specific window in development. Consistent with this, we saw very few myelin defects when DomA exposure occurred at 4 dpf – a time point after nascent myelin has been established (see below).

### Extrinsic factors that may influence the critical window for DomA toxicity

In zebrafish, 4 dpf is a time period at which myelin sheaths are already established. The absence of a myelin phenotype following exposures at 4 dpf suggests that DomA, at least at the doses used here, may not disrupt already established sheaths but rather may perturb the initial formation of myelin sheaths. Time-lapse imaging of the initial stages of axon wrapping and nascent myelination (from 2.5-3 dpf), provides additional evidence that DomA affects the ability of oligodendrocytes to initially wrap axons and form elongated myelin sheaths (Fig. 8B).

In addition to the intrinsic sensitivity of developing oligodendrocytes, it is likely that the 2 dpf window of susceptibility is also influenced by extrinsic factors that affect the distribution and availability of DomA to the cells and tissues of interest. One process that may influence DomA availability in the central nervous system is the development of the blood-brain barrier (BBB) − a structure that separates the blood from the brain parenchyma.^51^ The BBB is composed of tight junctions between endothelial cells that seal the intercellular cleft and prevent the diffusion of water-soluble molecules.^52–54^ As the BBB forms between 3-10 dpf, it progressively excludes smaller molecules over time. Thus, DomA may be excluded from the central nervous system to a greater degree during developmental periods past 2 dpf as the BBB matures.

DomA may also be less accessible to cell targets of interest later in development due to relatively higher excretion rates as the kidney matures. DomA is primarily cleared from the plasma via the kidneys, and nephrectomies in rodent models increase the plasma half-life of DomA.^55–57^ In zebrafish, glomerular filtration begins at around 2 dpf, while full maturation of the kidney occurs by 4 dpf.^58,59^ Thus, DomA may be more readily cleared during periods in development after 2 dpf.

### Transcriptional changes suggest defects in axon and myelin structures

RNAseq analysis identified genes and pathways that were consistent with the imaging and behavioral data. DomA exposure down-regulated genes required for maintaining myelin structure, including myelin protein zero (*mpz*) and (*mbpa*), along with genes required for maintaining axonal structure (*nefla, neflab, nefma, nefmb)* (Fig. 10). Thus, it is possible that DomA may be primarily targeting axons, and that the myelination defects may be a secondary effect. Alternatively, DomA may perturb oligodendrocyte development and myelin wrapping, leading to later axonal dysfunction. Further work is underway to investigate the potential axonal targets of DomA toxicity and to assess the contribution of the axonal disruptions to the myelin sheath phenotypes that we characterized here.^60^

**Figure 10.**
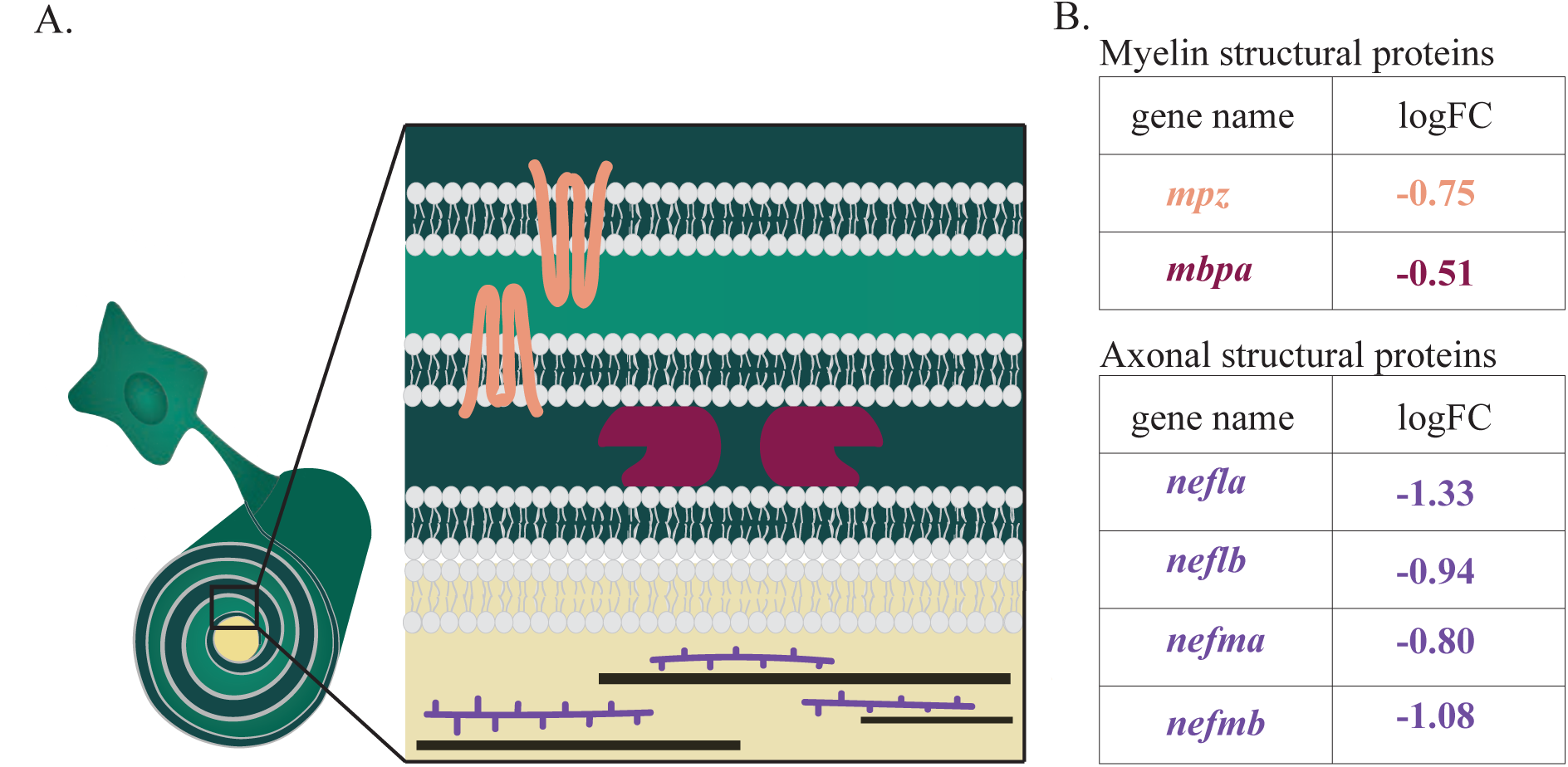
Domoic acid exposure at 2 dpf leads to reduced expression of key axonal and myelin structural proteins. (A) Schematic of the axon-myelin interface with a focus on selected myelin and axon structural proteins that are differentially expressed in DomA exposed fish. (B) Myelin and structural proteins that are differentially expressed with the log fold change (logFC). (-) indicates that the gene was down-regulated in DomA-exposed fish relative to controls.

RNAseq data show an increase in glial fibrillary acidic protein (*gfap*) expression following exposure to DomA. *gfap* is an intermediate filament protein whose upregulation in mammals is a hallmark of reactive gliosis – the response of glial cells following mechanical injury, excitotoxicity, or ischemia.^61,62^ In zebrafish, *gfap* expression is delayed following mechanical injury, and is expressed during the proliferation and recovery stages.^63–65^ The upregulation of *gfap* at 3 dpf suggests that exposure to DomA at 2 dpf may lead to injury and trigger repair mechanisms associated with increased *gfap* expression.

In addition, stathmin genes were overrepresented in our dataset. Stathmins destabilize microtubules by sequestering free tubulin. They are highly expressed in the developing nervous system and play critical roles in modulating neurite outgrowth and branching.^66,67^ It has been shown that the dysregulation of different stathmin genes (either through down- or up-regulation) can lead to alterations in microtubule density and axonal integrity.^67–69^

### Implications for human health

#### Timing and targets

This study provides a careful examination of potential windows of susceptibility to DomA exposure. The identification of key processes disrupted during these windows of susceptibility has important implications for identifying hazards for early developmental exposures in humans. Unlike in zebrafish, myelination in humans occurs over a prolonged period, starting *in utero* and continuing into early childhood and adolescence. The progression of myelination is mostly conserved across species, with myelination commencing in the periphery, brainstem, and spinal cord, then progressing rostrally to the forebrain.^70,71^ The most widespread and rapid period of myelination in humans occurs within the first two years of infancy.^72,73^ While most of the major tracts are myelinated by 3-5 years of age, myelination is now known to continue into adulthood, especially in cortical regions where changes in myelination are associated with experience and learning new skills.^74,75^ Thus, for humans, there may not be a single window of susceptibility, but rather multiple windows; domoic acid may perturb myelin formation in specific regions of the nervous system in which myelination coincides with the timing of exposures.

In this study, we showed that myelination was perturbed in the spinal cord – an understudied target tissue for domoic acid toxicity. Only one other study in rodents has investigated the spinal cord as a target tissue for DomA exposures. Wang et al. (2000) found that postnatal exposures to high doses of DomA led to spinal cord lesions by 2 hours post exposure, even in the absence of any histological damage to selected brain regions, including the well-known target, the hippocampus.^76^ Our study confirms the spinal cord as a potential target, and identifies myelination as the process perturbed in the spinal cord.

#### Behavioral analogies

We used startle response behavior as a functional readout of neurodevelopmental toxicity. Deficits in the kinematics of startle responses are reminiscent of motor deficits found in incidental human exposures, chronic exposures in primates, and developmental exposures in rodents. Adult humans acutely exposed to DomA developed sensorimotor neuropathy and axonopathy as assessed by electromyography.^77^ A subset of the primates exposed orally at or near the accepted daily tolerable dose of 0.075 mg/kg developed visible hand tremors.^78^ Rodents prenatally exposed to DomA (PND 10-17) developed aberrant gait patterns including impaired interlimb coordination and aberrant step sequence patterns.^21^

While there is evidence that DomA can perturb motor function, developmental exposures to DomA in rodents have not led to reductions in startle response amplitude during baseline conditions (prior to habituation or pre-pulse inhibition tests).^21,79–81^ This may be because exposures to DomA in these rodent models were done during a period that does not correspond to development of the startle circuit. Furthermore, there are some notable differences between rodent and fish startle, including distinct baseline startle kinematics and variations in the specific neuronal subsets in the circuits.^82–84^ Despite these differences, measuring startle response behavior in fish provides a tool to assess sensory processing and motor control and how these processes are perturbed by toxin exposure.

#### Doses and toxicokinetics

In all previous studies involving developmental exposure to DomA, ‘low doses’ have been defined based on the absence of acute neurotoxic symptoms, rather than by a specific dose. ‘Low doses’ are those that do not lead to classic acute symptoms that include tremors, scratching, and convulsions either in mothers (prenatal exposures) or in the pups directly exposed to DomA (postnatal exposures). While our study used nominal doses that were 3-to 260-fold lower than those used previously in zebrafish, these doses still led to transient neurotoxic effects in embryos. However, when directly comparing the weight-normalized amount of DomA, these doses are comparable to those used in the majority of the postnatal rodent studies.^11,23,79,80,85–88^ Assuming a 1.4 mg wet weight per embryo,^36^ the dosages at which embryos consistently exhibited myelin defects and behavioral deficits were 0.06-0.10 mg/kg DomA. In comparison, rodents who showed behavioral deficits following postnatal exposure were dosed subcutaneously with 7 injections of 0.005 and 0.020 mg/kg DomA between PND 8-14, leading to a comparable cumulative DomA dosage of 0.035-0.14 mg/kg.

The main challenge for translating findings in animal models to humans is the dearth of human exposure and toxicokinetic data. Human exposures to DomA are only estimated from consumption data, average weights of adults, and measured DomA concentrations in shellfish. Furthermore, the toxicokinetic behavior of DomA in humans is not well known. However, work in nonhuman primates shows that oral exposures to DomA lead to extended half-lives (almost 10x the length of the half-life following intravenous exposures).^89^ Furthermore, chronic exposure at or near the recognized tolerable daily intake level (0.075 and 0.150 mg/kg) leads to persistent hand tremors and disruptions to whole-brain connectivity.^78^

Even less information exists about the elimination and distribution in DomA in fetuses when mothers are exposed to DomA. One study in rodents showed that at one hour following intravenous injection of Dom A at GD13, the same concentrations of DomA were found in fetal brains, amniotic fluid, and maternal brains.^13^ This suggests that earlier in development there are no barriers for DomA entry to the fetal brain and that DomA in the fetal brain reaches equilibrium concentrations with DomA in the amniotic fluid. Emerging evidence from marine mammals shows that DomA can remain in the fetal fluids (amniotic and allantoic fluids) over prolonged periods of time.^16,17^ Thus, DomA may be recirculated within the fetal fluid compartments, allowing for continuous exposures in fetuses, even when maternal plasma has reached undetectable levels of DomA.

## CONCLUSIONS

DomA is a well-known developmental neurotoxin. However, few studies have been able to identify the cellular and molecular processes that underlie the observed behavioral deficits seen following developmental exposures. Using zebrafish, we were able to deliver DomA at specific developmental times and link behavioral deficits to structural changes in the neural circuit required for the behavior. The results from this study show that there is a critical window of susceptibility to DomA, and that exposure leads to altered expression of key axonal and myelin structural genes, disruptions to myelination, and later perturbations to startle behavior. These results establish the zebrafish as a model for investigating the cellular and molecular mechanisms underlying DomA-induced developmental neurotoxicity.

## MATERIALS AND METHODS

### Fish husbandry and lines used

These studies were approved by the Woods Hole Oceanographic Institution Animal Care and Use Committee (Assurance D16-00381 from the NIH Office of Laboratory Animal Welfare). Fish were maintained in recirculating tank systems that were specifically designed for zebrafish culture (Aquatic Habitats Inc., Apopka, FL). Temperature, lighting, and water quality were monitored daily and maintained according to recommendations from the Zebrafish International Resource Center. Fish were fed twice daily, once with live brine shrimp and once with the pellet feed Gemma Micro 300 (Skretting Inc., Tooele, UT). The afternoon before breeding, males and females were separated with a divider. The morning of the breeding, dividers were removed, and embryo collectors – containers with mesh on the top that let embryos filter to a catch basin – were placed in tanks with multiple breeding pairs for batch breeding unless otherwise noted. Embryos were collected and placed in petri dishes or in individual wells in a multi-well plate with 0.3x Danieau’s medium. Embryos were maintained at 28.5°C with a 14:10 light dark cycle during the experimental period.

The transgenic line *Tg(mbp:EGFP-CAAX)*^35^ in the AB background was used for behavioral, RNAseq, and myelin labeling experiments, while the double transgenic, *Tg(nkx2.2a:mEGFP)*,^90^ *Tg(sox10:RFP)*,^91^ was used for time lapse microscopy experiments.

### Domoic acid exposure paradigm

An initial pilot study was performed in which zebrafish embryos were exposed to DomA solutions (5-40 μM waterborne exposures). The absence of expected acute neurotoxicity even at high concentrations (data not shown) raised questions about whether DomA was being taken up by the embryos. Because of this, and to more precisely control the timing of exposure, we decided to use microinjection as the route of exposure.

Domoic acid was obtained in a 5 mg vial from Sigma-Aldrich, MO (D6152), and dissolved directly in the vial with diluted embryo medium (0.2x Danieau’s) to obtain a 20 mM solution. This was immediately used to generate stock concentrations of 0.675 μg/μl and 1.4 μg/μl. Aliquots (10 uL each) were stored at −20°C. Experiments were completed within 16 months of generating the stock. Working solutions were prepared fresh prior to microinjection by diluting the stock to obtain the appropriate doses. Microinjection needles were created from glass capillary tubes (058 mm inner diameter; World Precision Instruments, 1B100F-4) using a pipette puller (Sutter instrument model p-30, heat 750, pull= 0). Microinjections were performed using a Narishige IM-300 microinjector. The microinjector was calibrated to deliver 0.2 nL by adjusting the time (milliseconds) and pressure.

To determine the window of susceptibility for exposure at lower doses, DomA (0.09, 0.13, 0.14, 0.18 ng nominal dose) was intravenously microinjected into the common posterior cardinal vein at different developmental stages.^92^ Controls from the same breeding cohort were injected with the saline vehicle (0.2x Danieau’s). Supplemental Table 1 includes the developmental time ranges for each injection category. To perform intravenous microinjections, fish were dechorionated, anesthetized with tricaine methanesulfonate (MS222) (0.16%) then placed laterally on dishes coated with 1.5% agarose. An injection was deemed successful if there was a visible displacement of blood cells. Following injections, zebrafish were placed back in clean embryo media and monitored daily.

### Assessment of gross morphological defects and acute neurological phenotypes

Subsets of fish were imaged using brightfield microscopy to visualize potential gross morphological defects. The presence or absence of the swim bladder was scored blindly and then percentage was quantified for fish exposed to DomA at different doses and during different developmental stages. Images were white balance-corrected using Adobe Photoshop.

In a subset of the experiments, fish were kept individually in 48-well plates for phenotypic observation. Any mortalities, presence or absence of convulsions, pectoral flapping, and touch responses were recorded daily from the day after exposure to 5 dpf. Larvae were considered convulsing when whole body contractions were observed. Pectoral fin flapping was scored when larvae continued to rapidly move pectoral fins even when the fish were not actively swimming or attempting to right themselves. Touch responses were assessed using a tactile stimulus produced by an ‘embryo poker’ – a piece of fishing line (0.41 mm diameter) glued to a glass pipette tip. Larvae were identified as having no touch response when they were unable to perform body bends and swim away following tactile stimulation.

### Modeling the prevalence of neurotoxic phenotypes by dose, day of exposure, and day of observation

Following daily observation, generalized estimating equations (GEE) were used to model the effects of both DomA dose (as a continuous factor) and the number of days post-exposure (categorical factor) on the presence of acute neurological phenotypes (convulsions, pectoral flapping, and the lack of touch responses) (gee(), geepack R package).^49^ Observations of the same fish over multiple days were treated as repeated measures and were clustered by the “id” term. Separate GEE models were created for exposure to DomA at two developmental periods (1 and 2 dpf).

There were only single observations for fish exposed at 4 dpf (observed at 5 dpf). To determine whether DomA dose alters the presence of neurotoxic phenotypes one day post-exposure, a generalized linear model was formulated containing the different doses as predictors, and the presence of phenotypes as the response. To account for variability amongst trials, dispersion was estimated using the quasibinomial link function rather than the binomial one.

### Startle behavior set-up

The custom-built startle behavior set-up is shown in Fig 1B. The system includes a speaker (Visaton BG20-8 8” Full-Range Speaker with Whizzer Cone, #292-548) connected to an amplifier (100W TDA7498 Class-D Amplifier Board, #320-303) which serves as a source of auditory/vibrational stimuli. A hollow cylinder with a flat base was 3D printed and glued to the center of the speaker. This served as a platform to rest the plate that contained the fish (radius= 50 mm, height = 50mm). A 16-well acrylic plate (4.83 × 4.83 cm) was then designed to contain 16 larvae individually. This plate was based on a design from Wolman et al. (2011) that was comprised of laser cut acrylic pieces that were fused together using acrylic cement (Weld-On #3; IPS).^94^

The intensity and frequency of the auditory/vibrational stimuli were controlled using a pulse generator (PulsePal, Sansworks). Stimuli were coded to deliver 3 millisecond pulses of 1000 Hz frequency.

Groups of 16 larvae (7 dpf) were given 7 identical stimuli (41 dB) that were spaced 20 seconds apart to prevent habituation.^94^ A high-speed video camera (Edgertronic) was set at a 10% pre-trigger rate to capture 13 frames prior to the stimulus being elicited, while recording larval movements at 1000 frames per second.

### Measuring startle vibration

Vibration was measured using a 3-axis accelerometer (PCB Piezotronics, model W356B11). The output signal was first conditioned (PCB Piezotronics, Model 480B31) then passed through a dual channel analog filter (Model 3382, Krohn-Hite Corporation) using a 10 kHz low-pass cutoff frequency and 30 dB gain. Finally, the signal was collected by a data acquisition board (National Instruments Data Acquisition board, Model USB-6251). Raw voltage data were converted into acceleration units (m/s^2^) using manufacturer sensitivity values for each axis of the accelerometer. The Euclidian norm (vector sum) for the three acceleration signals was calculated to get the total acceleration. Individual peaks were identified, and metrics were calculated for the time window between 9 milliseconds prior to the peak to 50 ms after. The maximum value (peak) during each time window was taken as the zero to peak acceleration value for a given impulse, and this value was converted to dB using the following equation:

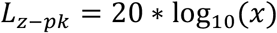

Where *L*_*z-pk*_ is the zero-to-peak acceleration level in dB re 1 m/s^2^, and *x* is the maximum acceleration level (of the Euclidian norm) over the peak analysis window.

### Startle behavioral analysis

High speed videos were converted into jpegs (.mov files with a minimal resolution of 720×720, 1/1008 shutter speed and a frame rate of 1000 frames/second). To reduce the noise and tracking errors, the background was subtracted, and the image contrast was enhanced using a custom script in MATLAB. FLOTE software^95^ was then used to analyze the jpegs. Quantitative attributes of the startle response measured include startle responsiveness (whether larvae responded or not), latency (delay time prior to startle), maximal bend angle, and maximal angular velocity during startle. The identities of individual larvae across the multiple stimuli were distinguished based on their position on a grid.

### Statistical modeling of startle responsiveness

Every fish was given 7 replicate auditory/vibrational stimuli, spaced 20 milliseconds apart. For all instances where a fish was successfully tracked, response rates were recorded. Percent response rates for individual fish were calculated (% responsiveness = number of times the fish responded / number of successfully tracked videos with a maximum of 7 tracks per individual fish). A mixed effects logistic regression model was used to identify treatment differences in percent responsiveness, with dose as a fixed effect and the replicate stimuli as a random effect using the ‘glmer’ function of lme4 package in R.^96^ A Dunnett post-hoc test was used to identify potential treatment differences in responsiveness (glht(), multcomp R package).^97^

### Identifying SLC versus LLC responses using mixture models

For all the fish that did respond, their startle responses were classified as either short latency c-bends (SLCs) or long latency c-bends (LLCs) based on an empirically determined latency cut-off. Latency cut-offs have been known to vary based on environmental conditions such as temperature.^95^ To empirically determine the cut-offs, clustering was done using a Gaussian mixture model, which fits two Gaussian distributions, and assigns each latency data point a probability of belonging to either of the two distributions (R package, mixtools).^98^ The cut-off for assigning a response as a SLC was 13 milliseconds – the latency with a greater than 50% probability of belonging to the first fitted Gaussian distribution (Supplemental Fig. 2). Startle responses that had latencies greater than 13 milliseconds were classified as LLCs.

### Analysis of treatment differences in startle response kinematics

There were several instances when individual fish performed a combination of LLC and SLC responses over the 7 replicate stimuli. For fish that did respond, their startle responses were classified as either SLCs (≤ 13 milliseconds) or LLCs (> 13 milliseconds). Kinematic responses from the two types of startle responses (SLC v. LLC) were analyzed separately based on previous research that shows they are driven by different neural circuits and have distinct kinematic characteristics.^95,99,100^ Following this classification, the median response of individual fish for each startle type was then used to identity treatment-specific differences in kinematics.

We first checked for normality and variance homogeneity in the data being analyzed. We used the Bartlett test to test for homogeneity in variances (bartlett.test(), R), and the Shapiro-Wilk’s method to test for normality (shapiro.test(), R). Kinematic data (bend angle, maximum angular velocity) showed departures from normality and had unequal variances. To account for this, we used nonparametric tests to determine whether fish exposed to various doses of DomA at different developmental periods had altered bend angles and maximal angular velocities.

Kinematic data from fish exposed to DomA at different development days were analyzed separately. For trials that contained a single dose of DomA, nonparametric Behrens-Fisher t-tests were used to test the alternative hypothesis that kinematics of fish exposed to DomA were different from their control counterparts (npar.t.test(), nparcomp package, R).^101^ With trials that contained multiple doses, nonparametric analyses with Dunnett-type intervals were done to compare each of the doses to the control (nparcomp(), nonparam package, R).^101^

Functions in the nparcomp package estimate the relative effects, which range from 0 to 1. Under the null hypothesis, the relative effect size is 0.5 – which represents a 50% probability (an equal probability) that the treated fish has a value greater than the control fish. The closer the estimated relative effect is to 1, the higher the probability that the measured kinematics in the treated group has a larger value than the control. In contrast, the closer the estimated relative effect is to 0, the higher the probability that the measured kinematic parameter in the treated group has a smaller value than the control.

### Startle kinematic analysis for interaction effects between dose and day of exposure

We then directly tested whether exposures that occurred on distinct developmental days influenced startle kinematics differently – in other words, if there is an interaction between dose and day injected. To examine this, we analyzed the subset of trials that had fish that were collected from the same breeding cohort at day 0 and then exposed to DomA at different developmental days (1, 2, or 4 dpf). Aligned Ranked Transformed ANOVA tests were done to determine whether there was an interaction between dose (0 versus 0.09 ng, or in a separate analysis, 0 versus 0.13 ng DomA) and day of exposure (1, 2, or 4 dpf) on startle kinematics (art(), ARTool R package).^102^ Difference-of-difference contrasts were then done to determine whether day of exposure affected treatment differences in kinematics (testContrasts(), Phia R package).^103^ Through this, we addressed questions such as, “is the difference in kinematics between control fish and those exposed at 2 dpf significant compared to the difference in kinematics between DomA and control fish when they are exposed at 1 or 4 dpf?”

### Fluorescence microscopy

Larvae were anesthetized in tricaine methanesulfonate (MS222) (0.16%), mounted laterally, and then imaged using either widefield epifluorescence microscopy or confocal microscopy. For images collected on the confocal microscope, fish were anesthetized and mounted laterally in 1.5% low melt agarose within glass bottom microscopy dishes (Nunc Glass bottom dishes 27mm). ‘Embryo pokers’ were used to orient the embryos onto their sides. Once the embryos were oriented correctly, the agarose was allowed to harden, and the microscopy dish was flooded with MS222. Fish were then imaged using the confocal microscope (Zeiss LSM-710 and LSM-780) with the 40x water objective (Zeiss C-Apochromat, NA= 1.1). Images were taken along the anterior spinal cord in the region around the 5^th^ and 10^th^ somites.

For images collected on the widefield epifluorescence microscope, a subset of fish were laterally mounted using 1.5% agarose. To allow for more rapid imaging of larvae, most larvae were oriented into custom-made acrylic molds that contained narrow channels where anesthetized larvae were positioned laterally using the embryo poker. Fish were imaged using the Zeiss inverted epifluorescence microscope with either a 20x (Fluar, NA = 0.75) or 10x (Fluar, NA = 0.5) objective. Images were taken along the anterior to medial spinal cord between somites 5-15.

### Analysis of the prevalence and severity of myelin phenotypes by dose and day of exposure

*Tg(mbp:EGFP-CAAX)* is a stable line in which EGFP is localized to cell membranes including myelin sheaths. We exposed *Tg(mbp:EGFP-CAAX)* fish to different doses of DomA at select developmental times and then imaged their spinal cords. Images were classified qualitatively into categories 0 through 5 based on severity in the myelin defect observed (Supplemental Fig. 4). Multinomial regression was used to model the effect of both dose and day injected on the distribution of the myelin severity phenotypes (multinom(), nnet R package).^104^

The overall significance of the dose and development day of exposures was obtained by performing an Analysis of Variance (ANOVA) on pairs of multinomial logistic regression models. The initial multinomial logistic regression model only included the dose of DomA as a predictor of the distribution of myelin phenotypes: β_0_ + β_dose_. The alternative model incorporated day of exposure: β_0_ + β_dose_ + β_DayExposure_. An ANOVA test was then used to determine whether the more complex alternative model was significantly better at capturing the data than the initial simpler one. A significant ANOVA result would determine whether day of exposure influences the distribution of the myelin phenotypes (anova(initial model, first alternative model), car package, R).^105^

Multinomial models were constructed to identify the effects of increasing doses of DomA on the distribution of these myelin phenotypes. To accomplish this, we used imaging data from fish exposed to varying doses of DomA at 2 dpf.

### Time-lapse microscopy

Embryos were exposed to DomA at 2 dpf, anesthetized, and mounted in 1.5% low melt agarose at around 2.25 dpf. Images were acquired on the LSM710 using the 20x dry (Plan-Apochromat 20x/0.8) objective. Z-stacks were acquired every 13-17 minutes over the course of 12-13 hours. For each embryo observed, maximum intensity projections of the z-stacks were then generated and compiled over time to generate the movie file (ZEN blue, ZEN black imaging software, Zeiss Microscopy).

### Experimental design for RNASeq

Three individual breeding tanks were set up with two males and one female per tank. Embryos collected from each tank were split so that some were injected with DomA (0.14 ng) and others with the saline vehicle control. Embryos were exposed to either the saline vehicle or to 0.14 ng of DomA at 2 dpf (between 48.5-51 hpf), then placed into 48-well plates for daily observation. Pools of 6 embryos from each of the three breeding sets were collected for RNAsequencing (n=3 per treatment) at 3 dpf (76 hpf). The remaining fish were used for imaging myelin at 5 dpf and for assessing startle behavior at 7 dpf (see below). At the end of the behavioral trial, a subset of the fish was snap frozen at 7 dpf (124 hpf) for RNA sequencing.

To ensure effectiveness of the exposure, a subset of exposed fish were imaged to visualize myelin structure at 5 dpf and then subjected to behavioral tests (startle response) at 7 dpf. Consistent with other experimental trials, there were differences in behavior and myelin labeling between DomA-exposed fish and controls (Supplemental Fig. 5). Fish exposed to DomA at 2 dpf had shorter bend angles and slower angular velocities relative to controls (Supplemental Fig. 5A and B). Also consistent with other experimental trials, only DomA-exposed larvae showed any visible myelin defects, with most of the fish having myelin defects that were in the second to highest severity (Category 3 = 21/ 49, Supplemental Fig. 5C). Phenotypic analysis thus validated the use of RNAseq to identify potential transcriptional changes from exposures.

### RNA Isolation and sequencing

RNA was isolated using the Zymo Direct-Zol kit (Catlog # R2062) and quantified using Nanodrop spectrophotometer. RNA quality was checked using the Bioanalyzer (Agilent technologies, CA) at the Harvard Biopolymers Facility, Cambridge, MA. RNA integrity number (RIN) of the samples was 8.2 or higher. Library preparation for single stranded RNAseq was done using the Illumina TruSeq total RNA library kit. Single-end 50 base pair sequencing was done on Illumina HiSeq2000 platform. Both library preparation and sequencing was performed at the Tufts University Core Facility (Boston, MA). Raw data files were assessed for quality using FastQC.^106^ Adapter trimming was done using Trimmomatic.^107^ Trimmed reads were aligned to the genome (GRCz10, version 84) using STAR aligner.^107,108^ HTSeq-count was used to count the number of reads mapped to the annotated regions of the genome.^109^ Differential gene expression (DGE) analysis was done using Bioconductor package, edgeR, following the DGE analysis pipeline outlined by Chen et al 2016.^110,111^ Raw and processed data files were deposited in NCBI Gene Expression Omnibus database (Accession number # GSE140045).

DGE analysis involved filtering genes with read counts less than 10/n, where n is the minimal library size, and then normalizing read counts. Negative binomial models were used to account for gene-specific variability from biological and technical sources. Multi-dimensional scaling plots were used to visualize the leading fold-changes (largest 500 log_2_ fold changes) between pairs of samples. False discovery rate of 5% (Benjamini-Hochberg method) was used as a statistical cutoff for identifying differentially expressed genes. Gene annotation was done using BioMart with the latest genome (GRCz11), and only annotated genes were used in pathway analysis. gProfiler was then used to identify enriched Gene Ontology (GO) terms and human phenology phenotypes.^112^ GO terms with evidence only from *in silico* curation methods were excluded from the enrichment analysis and a statistical significance level of less than or equal to 0.05 (adjusted p-value) was used.

## Supporting information

Collated supplemental data

## ACKNOWLEDGEMENTS

We would like to thank: Hanny E. Rivera and Andrew R. Solow for their advice on the statistical analysis, Harold Burgess for providing the FLOTE software for startle kinematic analysis, Ian T. Jones for measuring the vibrational output of the startle set-up, Benjamin G. Merrick for his help designing and building the startle apparatus, Louis Kerr for microscopy training and advice (MBL microscopy facility), and the labs who generously provided us with zebrafish transgenic lines to make this work possible − Bruce Appel (University of Colorado, Denver) and David Lyons (University of Edinburg).

## REFERENCES

1. Hampson DR, Huang X, Wells JW, Walter JA, Wright JLC. Interaction of domoic acid and several derivatives with kainic acid and AMPA binding sites in rat brain. Eur J Pharmacol. 1992;218(1):1–8. doi:10.1016/0014-2999(92)90140-Y

2. Lefebvre KA, Robertson A. Domoic acid and human exposure risks: a review. Toxicon. 2010;56(2):218–230. doi:10.1016/j.toxicon.2009.05.034

3. Perl TM, Bédard L, Kosatsky T, Hockin JC, Todd EC, Remis RS. An outbreak of toxic encephalopathy caused by eating mussels contaminated with domoic acid. N Engl J Med. 1990;322(25):1775–1780. doi:10.1056/NEJM199006213222504

4. Jeffery B, Barlow T, Moizer K, Paul S, Boyle C. Amnesic shellfish poison. Food Chem Toxicol. 2004;42(4):545–557. doi:10.1016/j.fct.2003.11.010

5. Wekell JC, Jurst J, Lefebvre KA. The origin of the regulatory limits for PSP and ASP toxins in shellfish. J Shellfish Res. 2010;23(July):927–930.

6. Mariën K. Establishing tolerable dungeness crab (Cancer magister) and razor clam (Siliqua patula) domoic acid contaminant levels. Environ Health Perspect. 1996;104(11):1230–1236. doi:10.1289/ehp.104-1469507

7. Grandjean P, Landrigan PJ. Neurobehavioural effects of developmental toxicity. Lancet Neurol. 2014;13(3):330–338. doi:10.1016/S1474-4422(13)70278-3

8. Andersen HR, Nielsen JB, Grandjean P. Toxicologic evidence of developmental neurotoxicity of environmental chemicals. Toxicology. 2000;144(1-3):121–127. doi:10.1016/S0300-483X(99)00198-5

9. Costa LG, Giordano G, Faustman EM. Domoic acid as a developmental neurotoxin. Neurotoxicology. 2010;31(5):409–423. doi:10.1016/j.neuro.2010.05.003

10. Tryphonas L, Truelove J, Nera E, Iverson F. Acute Neurotoxicity of Domoic Acid in the Rat. Toxicol Pathol. 1990;18(1):1–9. doi:10.1177/019262339001800101

11. Doucette TA, Bernard PB, Husum H, Perry MA, Ryan CL, Tasker RA. Low doses of domoic acid during postnatal development produce permanent changes in rat behaviour and hippocampal morphology. Neurotox Res. 2004;6(7-8):555–563. doi:10.1007/BF03033451

12. Xi D, Peng YG, Ramsdell JS. Domoic acid is a potent neurotoxin to neonatal rats. Nat Toxins. 1997;5(2):74–79. doi:10.1002/(SICI)(1997)5:2<74::AID-NT4>3.0.CO;2-I

13. Maucher JM, Ramsdell JS. Maternal-fetal transfer of domoic acid in rats at two gestational time points. Environ Health Perspect. 2007;115(12):1743–1746. doi:10.1289/ehp.10446

14. Ahrens MB, Li JM, Orger MB, et al. Brain-wide neuronal dynamics during motor adaptation in zebrafish. Nature. 2012;485(7399):471–477. doi:10.1038/nature11057

15. Scholin CA, Gulland F, Doucette GJ, et al. Mortality of sea lions along the central California coast linked to a toxic diatom bloom. Nature. 2000;403(6765):80–84. doi:10.1038/47481

16. Brodie Frances M D Gulland Denise J Greig EC, Hunter M, Jaakola J, Leger JS, Leighfield Frances M Van Dolah TA. Domoic acid causes reproductive failure in california sea lions (Zalophus Californianus). Mar Mammal Sci. 22AD;3(700-707). doi:10.1111/j.1748-7692.2006.00045.x

17. Lefebvre KA, Hendrix A, Halaska B, et al. Domoic acid in California sea lion fetal fluids indicates continuous exposure to a neuroteratogen poses risks to mammals. Harmful Algae. July 2018. doi:10.1016/J.HAL.2018.06.003

18. Rust L, Gulland F, Frame E, Lefebvre K. Domoic acid in milk of free living California marine mammals indicates lactational exposure occurs. Mar Mammal Sci. 2014;30(3):1272–1278. doi:10.1111/mms.12117

19. Maucher JM, Ramsdell JS. Domoic acid transfer to milk: evaluation of a potential route of neonatal exposure. Environ Health Perspect. 2005;113(4):461–464. doi:10.1289/ehp.7649

20. Tanemura K, Igarashi K, Matsugami T-R, Aisaki K, Kitajima S, Kanno J. Intrauterine environment-genome interaction and Children’s development (2): Brain structure impairment and behavioral disturbance induced in male mice offspring by a single intraperitoneal administration of domoic acid (DA) to their dams. J Toxicol Sci. 2009;34:SP279–SP286. doi:10.2131/jts.34.SP279

21. Shiotani M, Cole TB, Hong S, et al. Neurobehavioral assessment of mice following repeated oral exposures to domoic acid during prenatal development. Neurotoxicol Teratol. 2017;64:8–19. doi:10.1016/J.NTT.2017.09.002

22. Levin ED, Pizarro K, Pang WG, Harrison J, Ramsdell JS. Persisting behavioral consequences of prenatal domoic acid exposure in rats. Neurotoxicol Teratol. 27(5):719–725. doi:10.1016/j.ntt.2005.06.017

23. Perry MA, Ryan CL, Tasker RA. Effects of low dose neonatal domoic acid administration on behavioural and physiological response to mild stress in adult rats. Physiol Behav. 2009;98(1-2):53–59. doi:10.1016/J.PHYSBEH.2009.04.009

24. Burt MA, Ryan CL, Doucette TA. Altered responses to novelty and drug reinforcement in adult rats treated neonatally with domoic acid. Physiol Behav. 2008;93(1-2):327–336. doi:10.1016/j.physbeh.2007.09.003

25. Burt MA, Ryan CL, Doucette TA. Low dose domoic acid in neonatal rats abolishes nicotine induced conditioned place preference during late adolescence. Amino Acids. 2008;35(1):247–249. doi:10.1007/s00726-007-0584-2

26. Howe K, Clark MD, Torroja CF, et al. The zebrafish reference genome sequence and its relationship to the human genome. Nature. 2013;496(7446):498–503. doi:10.1038/nature12111

27. Tropepe V, Sive HL. Can zebrafish be used as a model to study the neurodevelopmental causes of autism? Genes Brain Behav. 2003;2(5):268–281. http://www.ncbi.nlm.nih.gov/pubmed/14606692. Accessed May 21, 2015.

28. Sumbre G, de Polavieja GG. The world according to zebrafish: how neural circuits generate behavior. Front Neural Circuits. 2014;8:91. doi:10.3389/fncir.2014.00091

29. Higashijima S, Masino MA, Mandel G, Fetcho JR. Imaging neuronal activity during zebrafish behavior with a genetically encoded calcium indicator. J Neurophysiol. 2003;90(6):3986–3997. doi:10.1152/jn.00576.2003

30. Fetcho JR, Higashijima S-I. Optical and genetic approaches toward understanding neuronal circuits in zebrafish. Integr Comp Biol. 2004;44(1):57–70. doi:10.1093/icb/44.1.57

31. Guo S. Linking genes to brain, behavior and neurological diseases: what can we learn from zebrafish? Genes, Brain Behav. 2004;3(2):63–74. doi:10.1046/j.1601-183X.2003.00053.x

32. Arrenberg AB, Driever W. Integrating anatomy and function for zebrafish circuit analysis. Front Neural Circuits. 2013;7:74. doi:10.3389/fncir.2013.00074

33. Eddins D, Cerutti D, Williams P, Linney E, Levin ED. Zebrafish provide a sensitive model of persisting neurobehavioral effects of developmental chlorpyrifos exposure: comparison with nicotine and pilocarpine effects and relationship to dopamine deficits. Neurotoxicol Teratol. 2010;32(1):99–108. doi:10.1016/j.ntt.2009.02.005

34. Pogoda H-M, Sternheim N, Lyons DA, et al. A genetic screen identifies genes essential for development of myelinated axons in zebrafish. Dev Biol. 2006;298(1):118–131. doi:10.1016/j.ydbio.2006.06.021

35. Almeida RG, Czopka T, Ffrench-Constant C, Lyons DA. Individual axons regulate the myelinating potential of single oligodendrocytes in vivo. Development. 2011;138(20):4443–4450. doi:10.1242/dev.071001

36. Tiedeken JA, Ramsdell JS, Ramsdell AF. Developmental toxicity of domoic acid in zebrafish (Danio rerio). Neurotoxicol Teratol. 2005;27:711–717.

37. Tiedeken JA, Ramsdell JS. Embryonic exposure to domoic Acid increases the susceptibility of zebrafish larvae to the chemical convulsant pentylenetetrazole. Environ Health Perspect. 2007;115(11):1547–1552. doi:10.1289/ehp.10344

38. Kirby BB, Takada N, Latimer AJ, et al. In vivo time-lapse imaging shows dynamic oligodendrocyte progenitor behavior during zebrafish development. Nat Neurosci. 2006;9(12):1506–1511. doi:10.1038/nn1803

39. Brösamle C, Halpern ME. Characterization of myelination in the developing zebrafish. Glia. 2002;39(1):47–57. doi:10.1002/glia.10088

40. Kolodziejczyk K, Saab AS, Nave K-A, Attwell D. Why do oligodendrocyte lineage cells express glutamate receptors? F1000 Biol Rep. 2010;2:57. doi:10.3410/B2-57

41. Patneau DK, Wright PW, Winters C, Mayer ML, Gallo V. Glial cells of the oligodendrocyte lineage express both kainate- and AMPA-preferring subtypes of glutamate receptor. Neuron. 1994;12(2):357–371. doi:10.1016/0896-6273(94)90277-1

42. Alberdi E, Sánchez-Gómez MV, Marino A, Matute C. Ca(2+) influx through AMPA or kainate receptors alone is sufficient to initiate excitotoxicity in cultured oligodendrocytes. Neurobiol Dis. 2002;9(2):234–243. doi:10.1006/nbdi.2001.0457

43. Matute C, Domercq M, Sánchez-Gómez M-V. Glutamate-mediated glial injury: mechanisms and clinical importance. Glia. 2006;53(2):212–224. doi:10.1002/glia.20275

44. Rosenberg PA, Dai W, Gan XD, et al. Mature myelin basic protein-expressing oligodendrocytes are insensitive to kainate toxicity. J Neurosci Res. 2003;71(2):237–245. doi:10.1002/jnr.10472

45. Deng W, Rosenberg P a, Volpe JJ, Jensen FE. Calcium-permeable AMPA/kainate receptors mediate toxicity and preconditioning by oxygen-glucose deprivation in oligodendrocyte precursors. Proc Natl Acad Sci U S A. 2003;100(11):6801–6806. doi:10.1073/pnas.1136624100

46. Gallo V, Zhou J, McBain C, Wright P, Knutson P, Armstrong R. Oligodendrocyte progenitor cell proliferation and lineage progression are regulated by glutamate receptor-mediated K+ channel block. J Neurosci. 1996;16(8):2659–2670. http://www.jneurosci.org/content/16/8/2659.short. Accessed December 7, 2014.

47. Gudz TI, Komuro H, Macklin WB. Glutamate stimulates oligodendrocyte progenitor migration mediated via an alphav integrin/myelin proteolipid protein complex. J Neurosci. 2006;26(9):2458–2466. doi:10.1523/JNEUROSCI.4054-05.2006

48. Matute C. Characteristics of acute and chronic kainate excitotoxic damage to the optic nerve. Proc Natl Acad Sci U S A. 1998;95(17):10229–10234. doi:pnas.95.17.10229

49. Verity AN, Campagnoni AT. Myelination and Its Underlying Mechanisms Regional Expression of Myelin Protein Genes in the Developing Mouse Brain: In Situ Hybridization Studies. Vol 21.; 1988. https://onlinelibrary.wiley.com/doi/pdf/10.1002/jnr.490210216. Accessed February 8, 2019.

50. Foran DR, Peterson AC. Myelin acquisition in the central nervous system of the mouse revealed by an MBP-Lac Z transgene. J Neurosci. 1992;12(12):4890–4897. doi:10.1523/JNEUROSCI.12-12-04890.1992

51. Eliceiri BP, Gonzalez AM, Baird A. Zebrafish Model of the Blood-Brain Barrier: Morphological and Permeability Studies. In: Methods in Molecular Biology (Clifton, N.J.). Vol 686.; 2011:371–378. doi:10.1007/978-1-60761-938-3_18

52. Fleming A, Diekmann H, Goldsmith P. Functional Characterisation of the Maturation of the Blood-Brain Barrier in Larval Zebrafish. Del Bene F, ed. PLoS One. 2013;8(10):e77548. doi:10.1371/journal.pone.0077548

53. Jeong J-Y, Kwon H-B, Ahn J-C, et al. Functional and developmental analysis of the blood–brain barrier in zebrafish. Brain Res Bull. 2008;75(5):619–628. doi:10.1016/J.BRAINRESBULL.2007.10.043

54. Xie J, Farage E, Sugimoto M, Anand-Apte B. A novel transgenic zebrafish model for blood-brain and blood-retinal barrier development. BMC Dev Biol. 2010;10(1):76. doi:10.1186/1471-213X-10-76

55. Preston E, Hynie I. Transfer constants for blood-brain barrier permeation of the neuroexcitatory shellfish toxin, domoic acid. Can J Neurol Sci. 1991;18(1):39–44. http://www.ncbi.nlm.nih.gov/pubmed/2036614. Accessed September 17, 2018.

56. Suzuki CAM, Hierlihy SL. Renal clearance of domoic acid in the rat. Food Chem Toxicol. 1993;31(10):701–706. doi:10.1016/0278-6915(93)90140-T

57. Lefebvre KA, Noren DP, Schultz IR, Bogard SM, Wilson J, Eberhart BT. Uptake, tissue distribution and excretion of domoic acid after oral exposure in coho salmon (Oncorhynchus kisutch). Aquat Toxicol. 2007;81(3):266–274. doi:10.1016/j.aquatox.2006.12.009

58. Drummond IA, Davidson AJ. Zebrafish Kidney Development. Methods Cell Biol. 2010;100:233–260. doi:10.1016/B978-0-12-384892-5.00009-8

59. Drummond IA. Kidney Development and Disease in the Zebrafish. J Am Soc Nephrol. 2005;16:299–304. doi:10.1681/ASN.2004090754

60. Panlilio JM, Aluru N, Hahn ME. Early Developmental Exposure to Low Levels of Domoic Acid, a Harmful Algal Bloom Toxin, Disrupts Myelination, leading to Behavioral Effects. Toxicol Suppl to Toxicol Sci. 2019;168(1):Abstract #1691.

61. Burtrum D, Silverstein FS. Excitotoxic Injury Stimulates Glial Fibrillary Acidic Protein mRNA Expression in Perinatal Rat Brain. Exp Neurol. 1993;121(1):127–132. doi:10.1006/exnr.1993.1078

62. Nielsen AL, Jørgensen AL. Structural and functional characterization of the zebrafish gene for glial fibrillary acidic protein, GFAP. Gene. 2003;310:123–132. doi:10.1016/S0378-1119(03)00526-2

63. Lam CS, März M, Strähle U. gfap and nestin reporter lines reveal characteristics of neural progenitors in the adult zebrafish brain. Dev Dyn. 2009;238(2):475–486. doi:10.1002/dvdy.21853

64. Hui SP, Nag TC, Ghosh S. Characterization of Proliferating Neural Progenitors after Spinal Cord Injury in Adult Zebrafish. Thummel R, ed. PLoS One. 2015;10(12):e0143595. doi:10.1371/journal.pone.0143595

65. Grupp L, Wolburg H, Mack AF. Astroglial structures in the zebrafish brain. J Comp Neurol. 2010;518(21):4277–4287. doi:10.1002/cne.22481

66. Grenningloh G, Soehrman S, Bondallaz P, Ruchti E, Cadas H. Role of the microtubule destabilizing proteins SCG10 and stathmin in neuronal growth. J Neurobiol. 2004;58(1):60–69. doi:10.1002/neu.10279

67. Wen H-L, Lin Y-T, Ting C-H, Lin-Chao S, Li H, Hsieh-Li HM. Stathmin, a microtubule-destabilizing protein, is dysregulated in spinal muscular atrophy†. Hum Mol Genet. 2010;19(9):1766–1778. doi:10.1093/hmg/ddq058

68. Cheng HW, Jiang T, Mori N, McNeill TH. Upregulation of stathmin (p19) gene expression in adult rat brain during injury-induced synapse formation. Neuroreport. 1997;8(17):3691–3695. doi:10.1097/00001756-199712010-00007

69. Wen H-L, Ting C-H, Liu H-C, Li H, Lin-Chao S. Decreased stathmin expression ameliorates neuromuscular defects but fails to prolong survival in a mouse model of spinal muscular atrophy. Neurobiol Dis. 2013;52:94–103. doi:10.1016/J.NBD.2012.11.015

70. Rice D, Barone S. Critical periods of vulnerability for the developing nervous system: evidence from humans and animal models. Environ Health Perspect. 2000;108 Suppl:511–533. doi:10.1289/ehp.00108s3511

71. Tanaka S, Mito T, Takashima S. Progress of myelination in the human fetal spinal nerve roots, spinal cord and brainstem with myelin basic protein immunohistochemistry. Early Hum Dev. 1995;41(1):49–59. doi:10.1016/0378-3782(94)01608-R

72. Kinney HC, Volpe JJ. Myelination Events. In: Volpe’s Neurology of the Newborn. Elsevier; 2018:176–188. doi:10.1016/B978-0-323-42876-7.00008-9

73. Kinney HC, Ann brody B, Kloman AS, Gilles FH. Sequence of Central Nervous System Myelination in Human Infancy. II. Patterns of Myelination in Autopsied Infants. J Neuropathol Exp Neurol. 1988;47(3):217–234. doi:10.1097/00005072-198805000-00003

74. Fields RD. Myelination: an overlooked mechanism of synaptic plasticity? Neuroscientist. 2005;11(6):528–531. doi:10.1177/1073858405282304

75. Pajevic S, Basser PJ, Fields RD. Role of myelin plasticity in oscillations and synchrony of neuronal activity. Neuroscience. 2014;276:135–147. doi:10.1016/j.neuroscience.2013.11.007

76. Wang GJ, Schmued LC, Andrews AM, Scallet AC, Slikker W, Binienda Z. Systemic administration of domoic acid-induced spinal cord lesions in neonatal rats. J Spinal Cord Med. 2000;23(1):31–39. http://www.ncbi.nlm.nih.gov/pubmed/10752872. Accessed April 22, 2015.

77. Teitelbaum JS, Zatorre RJ, Carpenter S, et al. Neurologic Sequelae of Domoic Acid Intoxication Due to the Ingestion of Contaminated Mussels. N Engl J Med. 1990;322(25):1781–1787. doi:10.1056/NEJM199006213222505

78. Petroff R, Richards T, Crouthamel B, et al. Chronic, Low-Level Oral Exposure to Marine Toxin, Domoic Acid, Alters Whole Brain Morphometry in Nonhuman Primates. Neurotoxicology. 2019. doi:10.1101/439109

79. Adams AL, Doucette TA, Ryan CL. Altered pre-pulse inhibition in adult rats treated neonatally with domoic acid. Amino Acids. 2008;35(1):157–160. doi:10.1007/s00726-007-0603-3

80. Marriott AL, Ryan CL, Doucette TA. Neonatal domoic acid treatment produces alterations to prepulse inhibition and latent inhibition in adult rats. Pharmacol Biochem Behav. 2012;103(2):338–344. doi:10.1016/j.pbb.2012.08.022

81. Zuloaga DG, Lahvis GP, Mills B, Pearce HL, Turner J, Raber J. Fetal domoic acid exposure affects lateral amygdala neurons, diminishes social investigation and alters sensory-motor gating. Neurotoxicology. 2016;53:132–140. doi:10.1016/J.NEURO.2016.01.007

82. Koch M. The neurobiology of startle. Prog Neurobiol. 1999;59(2):107–128. http://www.ncbi.nlm.nih.gov/pubmed/10463792. Accessed May 1, 2015.

83. Eaton RC, Lee RKK, Foreman MB. The Mauthner cell and other identified neurons of the brainstem escape network of fish. Prog Neurobiol. 2001;63(4):467–485. doi:10.1016/S0301-0082(00)00047-2

84. Yeomans JS, Frankland PW. The acoustic startle reflex: neurons and connections. Brain Res Rev. 1995;21(3):301–314. doi:10.1016/0165-0173(96)00004-5

85. Bernard PB, MacDonald DS, Gill DA, Ryan CL, Tasker RA. Hippocampal mossy fiber sprouting and elevated trkB receptor expression following systemic administration of low dose domoic acid during neonatal development. Hippocampus. 2007;17(11):1121–1133. doi:10.1002/hipo.20342

86. Ryan CL. Hippocampal mossy fiber sprouting and elevated trkB receptor expression following systemic administration of low dose domoic acid during neonatal development. 2007. doi:10.1002/hipo.20342

87. Gill DA, Bastlund JF, Watson WP, Ryan CL, Reynolds DS, Tasker RA. Neonatal exposure to low-dose domoic acid lowers seizure threshold in adult rats. Neuroscience. 2010;169(4):1789–1799. doi:10.1016/j.neuroscience.2010.06.045.

88. Tasker RAR, Perry MA, Doucette TA, Ryan CL. NMDA receptor involvement in the effects of low dose domoic acid in neonatal rats. Amino Acids. 2005;28(2):193–196. doi:10.1007/s00726-005-0167-z

89. Jing J, Petroff R, Shum S, et al. Toxicokinetics and Physiologically Based Pharmacokinetic Modeling of the Shellfish Toxin Domoic Acid in Nonhuman Primates. Drug Metab Dispos. 2018;46:155–165. doi:10.1124/dmd.117.078485

90. Kucenas S, Snell H, Appel B. nkx2.2a promotes specification and differentiation of a myelinating subset of oligodendrocyte lineage cells in zebrafish. Neuron Glia Biol. 2008;4(2):71–81. doi:10.1017/S1740925X09990123

91. Takada N, Kucenas S, Appel B. Sox10 is necessary for oligodendrocyte survival following axon wrapping. Glia. 2010;58(8):996–1006. doi:10.1002/glia.20981

92. Cianciolo Cosentino C, Roman BL, Drummond IA, Hukriede NA. Intravenous microinjections of zebrafish larvae to study acute kidney injury. J Vis Exp. 2010;(42). doi:10.3791/2079

93. Halekoh U, Højsgaard S, Yan J. The *R* Package geepack for Generalized Estimating Equations. J Stat Softw. 2006;15(2):1–11. doi:10.18637/jss.v015.i02

94. Wolman MA, Jain RA, Liss L, Granato M. Chemical modulation of memory formation in larval zebrafish. Proc Natl Acad Sci U S A. 2011;108(37):15468–15473. doi:10.1073/pnas.1107156108

95. Burgess HA, Granato M. Modulation of locomotor activity in larval zebrafish during light adaptation. J Exp Biol. 2007;210(Pt 14):2526–2539. doi:10.1242/jeb.003939

96. Bates D, Mächler M, Bolker B, Walker S. Fitting Linear Mixed-Effects Models using lme4. June 2014. http://arxiv.org/abs/1406.5823. Accessed December 25, 2018.

97. Hothorn T, Bretz F, Westfall P. The Multcomp Package Title Simultaneous Inference for General Linear Hypotheses.; 2007. http://132.180.15.2/math/statlib/R/CRAN/doc/packages/multcomp.pdf. Accessed December 25, 2018.

98. Benaglia T, Chauveau D, Hunter D, Young D. mixtools: An R Package for Analyzing Finite Mixture Models. J Stat Softw. 2009;32(6):1–29. https://hal.archives-ouvertes.fr/hal-00384896/. Accessed December 25, 2018.

99. O’Malley DM, Kao YH, Fetcho JR. Imaging the functional organization of zebrafish hindbrain segments during escape behaviors. Neuron. 1996;17(6):1145–1155. http://www.ncbi.nlm.nih.gov/pubmed/8982162. Accessed January 29, 2015.

100. Marsden KC, Granato M. In Vivo Ca(2+) Imaging Reveals that Decreased Dendritic Excitability Drives Startle Habituation. Cell Rep. 2015;13(9):1733–1740. doi:10.1016/j.celrep.2015.10.060

101. Konietschke F, Placzek M, Schaarschmidt F, Hothorn LA. nparcomp : An *R* Software Package for Nonparametric Multiple Comparisons and Simultaneous Confidence Intervals. J Stat Softw. 2015;64(9). doi:10.18637/jss.v064.i09

102. Wobbrock JO, Findlater L, Gergle D, Higgins JJ. The aligned rank transform for nonparametric factorial analyses using only anova procedures. In: Proceedings of the 2011 Annual Conference on Human Factors in Computing Systems - CHI ‘11. New York, New York, USA: ACM Press; 2011:143. doi:10.1145/1978942.1978963

103. Rosario-Martinez H De, Fox J, Team RC. Post-Hoc Interaction Analysis [R package phia version 0.2-1]. https://cran.r-project.org/web/packages/phia/index.html. Accessed December 25, 2018.

104. Ripley B, Venables W. Package “Nnet.”; 2016. http://www.stats.ox.ac.uk/pub/MASS4/. Accessed February 7, 2019.

105. Fox J, Weisberg S, Price B, et al. Package “car.” In: An R Companion to Applied Regression. SAGE Publications; 2018. ftp://ftp.math.ethz.ch/sfs/pub/Software/CRAN/web/packages/car/car.pdf. Accessed February 7, 2019.

106. Andrews S. FastQC: a quality control tool for high throughput sequence data.

107. Bolger AM, Lohse M, Usadel B. Trimmomatic: a flexible trimmer for Illumina sequence data. Bioinformatics. 2014;30(15):2114–2120. doi:10.1093/bioinformatics/btu170

108. Dobin A, Davis CA, Schlesinger F, et al. STAR: ultrafast universal RNA-seq aligner. Bioinformatics. 2013;29(1):15–21. doi:10.1093/bioinformatics/bts635

109. Anders S, Pyl PT, Huber W. HTSeq--a Python framework to work with high-throughput sequencing data. Bioinformatics. 2015;31(2):166–169. doi:10.1093/bioinformatics/btu638

110. Robinson MD, McCarthy DJ, Smyth GK. edgeR: a Bioconductor package for differential expression analysis of digital gene expression data. Bioinformatics. 2010;26(1):139–140. doi:10.1093/bioinformatics/btp616

111. Chen Y, Lun ATL, Smyth GK. From reads to genes to pathways: differential expression analysis of RNA-Seq experiments using Rsubread and the edgeR quasi-likelihood pipeline. F1000Research. 2016;5:1438. doi:10.12688/f1000research.8987.2

112. Reimand J, Arak T, Adler P, et al. g:Profiler—a web server for functional interpretation of gene lists (2016 update). Nucleic Acids Res. 2016;44(W1):W83-W89. doi:10.1093/nar/gkw199

113. Zhang Y, Kecskés A, Copmans D, et al. Pharmacological characterization of an antisense knockdown zebrafish model of Dravet syndrome: Inhibition of epileptic seizures by the serotonin agonist fenfluramine. PLoS One. 2015;10(5). doi:10.1371/journal.pone.0125898

