## Supplementary material for "Domoic acid disruption of neurodevelopment and behavior involves altered myelination in the spinal cord": Collated supplemental data

### SUPPLEMENTAL RESULTS AND DISCUSSION

*Mortality and gross morphological effects of DomA*

When DomA (0.09-0.14 ng) was injected intravenously into *Tg(mbp:EGFP-CAAX)* embryos or larvae, there were no significant differences in mortality between DomA-exposed versus vehicle-treated (control) fish for any of the developmental exposure time points tested (1-4 dpf) (Supplemental Table 2).

A majority of the larvae exposed to DomA at 2 dpf did not have inflated swim bladders at 5 dpf (Supplemental Fig 1B, Supplemental Table 3), and a subset of them had a curved body axis. Phenotypes are consistent with other zebrafish models with neurodevelopmental abnormalities.[113]

Furthermore, some larvae injected at 4 dpf with the highest dose of DomA (0.18 ng) had brains with a darkened appearance. These 'opaque' brains were signs of widespread apoptosis or necrosis, suggesting that this dose could lead to widespread neurotoxicity (Supplemental Table 4). Based on these results, DomA doses of 0.14 ng or lower were primarily used to assess startle response behavior and gene expression analyses.

*Acute neurotoxicity of DomA*

Fish exposed to all doses of DomA (0.09-0.18 ng) at 1 dpf had significantly reduced touch responses as compared to controls (Supplemental Table 5 and 8, Supplemental Fig 1C). Furthermore, the prevalence of reduced touch responsiveness increased with dose of DomA ($p = 1.5e-8$). This phenotype was transient; the prevalence of touch response deficits dropped after the first day post-exposure ($p= 0.0028$).

Similarly, fish exposed to all doses of DomA at 2 dpf also had reduced touch responses, with a higher prevalence at high doses ($p= 1.5e-8$, Supplemental Table 6 and 8, Supplemental Fig 1C). However, unlike fish exposed at 1 dpf, those exposed to a middle dose (0.13 ng of DomA) had touch response deficits similar to those exposed to the highest dose (0.18 ng). Similar to 1 dpf, the touch response phenotype was transient, with significant reductions in touch response deficits after the first day post-exposure ($p= 6.6e-6$). At 4 dpf, only fish given the highest dose of DomA (0.18ng) exhibited any touch response deficits (Supplemental Table 7, Supplemental Fig 1C).

The presence of convulsions or pectoral fin flapping was also monitored (Supplemental Fig 1D and Supplemental Table 9). Fish exposed to DomA at both 1 and 2 dpf (but not at 4 dpf) had a dose-dependent increase in the prevalence of the phenotype ($p =0.0001$ for 1 dpf exposed fish, $p = 1.2e-7$ for 2 dpf exposed fish). There was a higher overall prevalence of this phenotype in 2-dpf exposed fish. Both touch response deficits and convulsions were transient, with the loss of convulsions and pectoral fin flapping by 2 days post exposure in 1 dpf injected fish (p= 0.001), and by 1 day post exposure in 2 dpf injected fish (p= 2.2e-9).

Overall, these results show that exposure to DomA (0.14 ng or lower) did not lead to appreciable mortality, but resulted in transient acute neurotoxic phenotypes that lasted one day after exposure. While fish exposed at both 1 and 2 dpf exhibited these neurotoxic phenotypes, fish exposed to DomA at 2 dpf had these phenotypes at a higher prevalence than those exposed at 1 dpf.

*Acute neurotoxic phenotypes are transient and dose- and exposure time-dependent*

Fish exposed to DomA at 1 and 2 dpf exhibited all the acute neurotoxic symptoms previously described in developmental exposure studies in zebrafish.[36] However, Tiedekken et al. (2005) found that high doses of DomA led to neurotoxic effects that occur during the larval stages.[36] In contrast, our experiments showed that the loss of a touch response and presence of pectoral fin flapping and tonic-clonic convulsions were transient, lasting only one to two days after exposure during the embryonic period (Supplemental Fig. 1C, 1D).

Both dose and timing of exposures significantly influenced the presence of acute neurotoxic symptoms. Fish exposed at 4 dpf had no significant acute neurotoxic symptoms, while fish exposed at both 1 and 2 dpf had touch response deficits. While exposures during both these developmental periods led to these neurotoxic symptoms, embryos exposed to DomA at 2 dpf were more sensitive and showed more pronounced touch response deficits than those exposed at 1 dpf (Supplemental Fig 1D). These results suggest that embryos were more sensitive to DomA at 2 dpf, exhibiting a higher prevalence in acute neurotoxic symptoms at the lower doses tested.

### SUPPLEMENTAL FIGURE LEGENDS

**Supplemental Figure 1: Domoic acid-exposed larvae have morphological and acute neurotoxic phenotypes that vary based on exposure time and dose.**

**(A1)** Representative brightfield image of control larvae with an inflated swim bladder.

**(A2)** Representative brightfield image of DomA-exposed larvae with an inflated swim bladder (teal), and without an inflated swim bladder (peach).

**(B)** Presence or absence of the inflated swim bladder by treatment, dose, and time of exposure.

**(C)** The percentage of embryos with touch response deficits were recorded one day post-exposure until 5 dpf. Fish were exposed to different doses of DomA (0.09 ng- 0.18 ng) at 1, 2, and 4 dpf. Lines are dodged along the x axis. ± standard errors of the mean between repeated experiments.

**(D)** The same fish population observed in Fig. 1C were also monitored for the presence of convulsions or pectoral fin flapping from one day post-exposure until 5 dpf.

**Supplemental figure 2: Startle behavioral classification**

Density histogram of the latency distribution for control fish. Overlaid are two Gaussian curves that were fit to the data. The teal curve represents the SLC distribution, and peach represents the LLC distribution. The solid black vertical line at 13 milliseconds represents the cut-off by which there is a greater than 50% probability of a given data point belonging to either modeled distribution.

**Supplemental Figure 3. Exposure to domoic acid at 2 dpf (but not 1 or 4 dpf) alters LLC startle response kinematics at the lowest dose.**

(**A**) Bend angles during LLC startles in fish exposed during different exposure days (1, 2 and 4 dpf) to the nominal dose of 0.09 ng of DomA and **(B)** to a dose that ranged from 0.126- 0.144 (labelled as 0.13ng of DomA). **(C)** Maximal angular velocity during LLC startles in fish exposed during different exposure days to 0.09 ng of DomA and to **(D)** 0.13ng of DomA. Each point represents the median kinematic response of an individual larvae to multiple identical stimuli.

* $p < 0.05$, **= $p < 0.01$, *** $p < 0.001$.

Figure supplement: Supplemental video 1is a sample startle response.

**Supplemental Figure 4: Qualitative myelin phenotype scoring**

Myelinated axonal tracks in the dorsal and ventral spinal cord. Fish were blindly classified into 6 categories (0-5) based on the severity in the myelin defect. Representative confocal and widefield fluorescence microscopy images are shown for each severity classification.

**Supplemental Figure 5: Startle kinematics and myelin sheath imaging in fish used for RNASeq**

(A) Startle bend angle for both DomA treated and control fish undergoing SLC or LLC startle responses. Each point represents the median kinematic response of individual larvae to multiple identical stimuli. (B) Maximal angular velocity obtained during the initial bend for both DomA and control fish undergoing SLC or LLC startle responses. (C) Distribution of myelin phenotypes in control (0 ng) and DomA (0.14 ng) exposed larvae at 2 dpf. Each color denotes a separate myelin category each fish was classified under. Numbers above denote the total number of fish per treatment group.

*** = $p < 0.0001$; Scale bar = 100 μm

**Supplemental Video 1: Acoustic startle response**

A 16 well plate containing control fish in the top two rows) and DomA exposed fish (2 dpf injected, 0.14ng) in the bottom two rows. Fish were subjected to a vibrational stimulus, while their responses were recorded at 1000 frames per second.

**Supplemental Video 2: Time-lapse of *Tg(sox10:RFP)* x *Tg(nkx2.2a:mEGFP)* control fish**

Time-lapse sequence of the spinal cord of control fish injected at 2 dpf taken from 2.5- 3 dpf. RFP labels cell bodies of cells from the oligodendrocyte lineage. mEGFP expression labels oligodendrocyte membrane processes that wrap axons and become elongated, nascent myelin sheaths.

**Supplemental Video 3: Time-lapse of *Tg(sox10:RFP)* x *Tg(nkx2.2a:mEGFP)* DomA exposed fish**

Time-lapse sequence of the spinal cord of DomA-exposed fish (0.14 ng) injected at 2 dpf taken from 2.5-3 dpf. RFP labels cell bodies of cells from the oligodendrocyte lineage. mEGFP expression labels oligodendrocyte membrane processes that form unusual circular membranes.

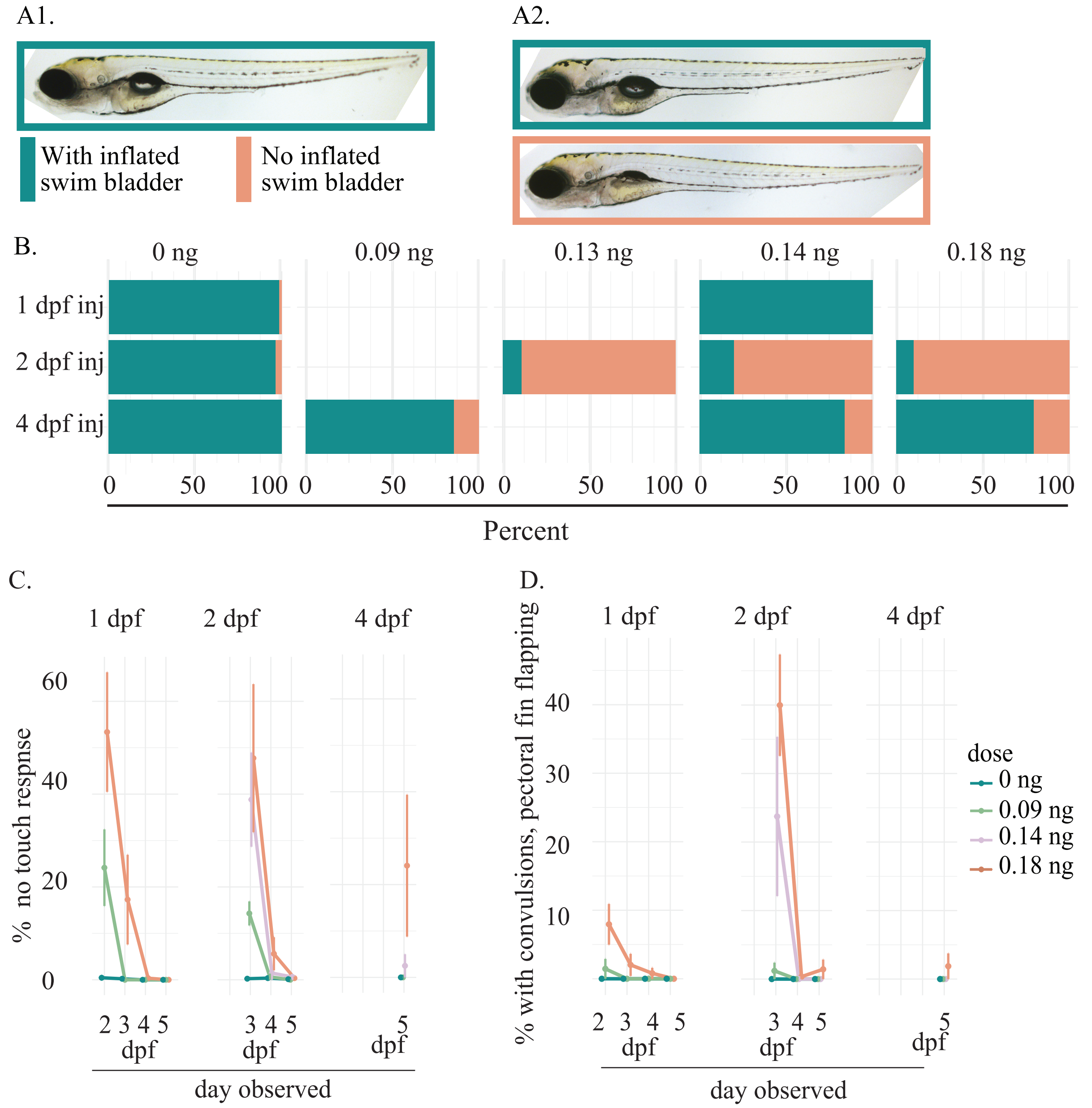


**Supplemental figure 1: Domoic acid-exposed larvae have morphological and acute neurotoxic phenotypes that vary based on exposure time and dose.**
**(A1)** Brightfield image of control larvae with an inflated swim bladder.
**(A2)** Brightfield image of DomA-exposed larvae with an inflated swim bladder (teal), and without an inflated swim bladder (peach).
**(B)** Presence or absence of the inflated swim bladder by treatment, dose, and time of exposure.
**(C)** The percentage of embryos with touch response deficits were recorded one day post-exposure until 5 dpf. Fish were exposed to different doses of DomA (0.09 ng- 0.18 ng) at 1, 2, and 4 dpf. Lines are dodged along the x axis. ± standard errors of the mean between repeated experiments.
**(D)** The same fish population observed in Fig. 1C were also monitored for the presence of convulsions or pectoral fin flapping from one day post-exposure until 5 dpf.

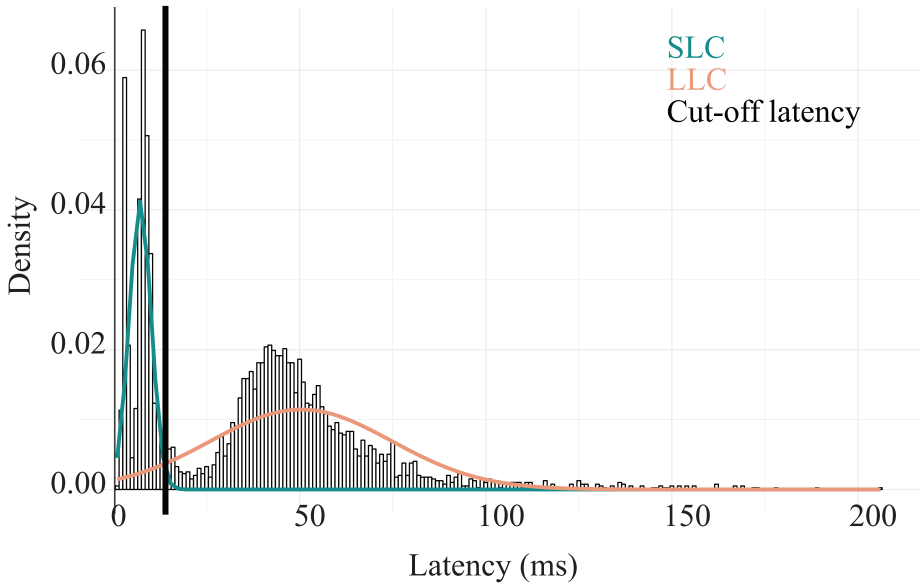


**Supplemental figure 2: Startle behavioral classification**
Density histogram of the latency distribution for control fish. Overlaid are two Gaussian curves that were fit to the data. The teal curve represents the SLC distribution, and peach represents the LLC distribution. The solid black vertical line at 13 milliseconds represents the cut-off by which there is a greater than 50% probability of a given data point belonging to either modeled distribution.

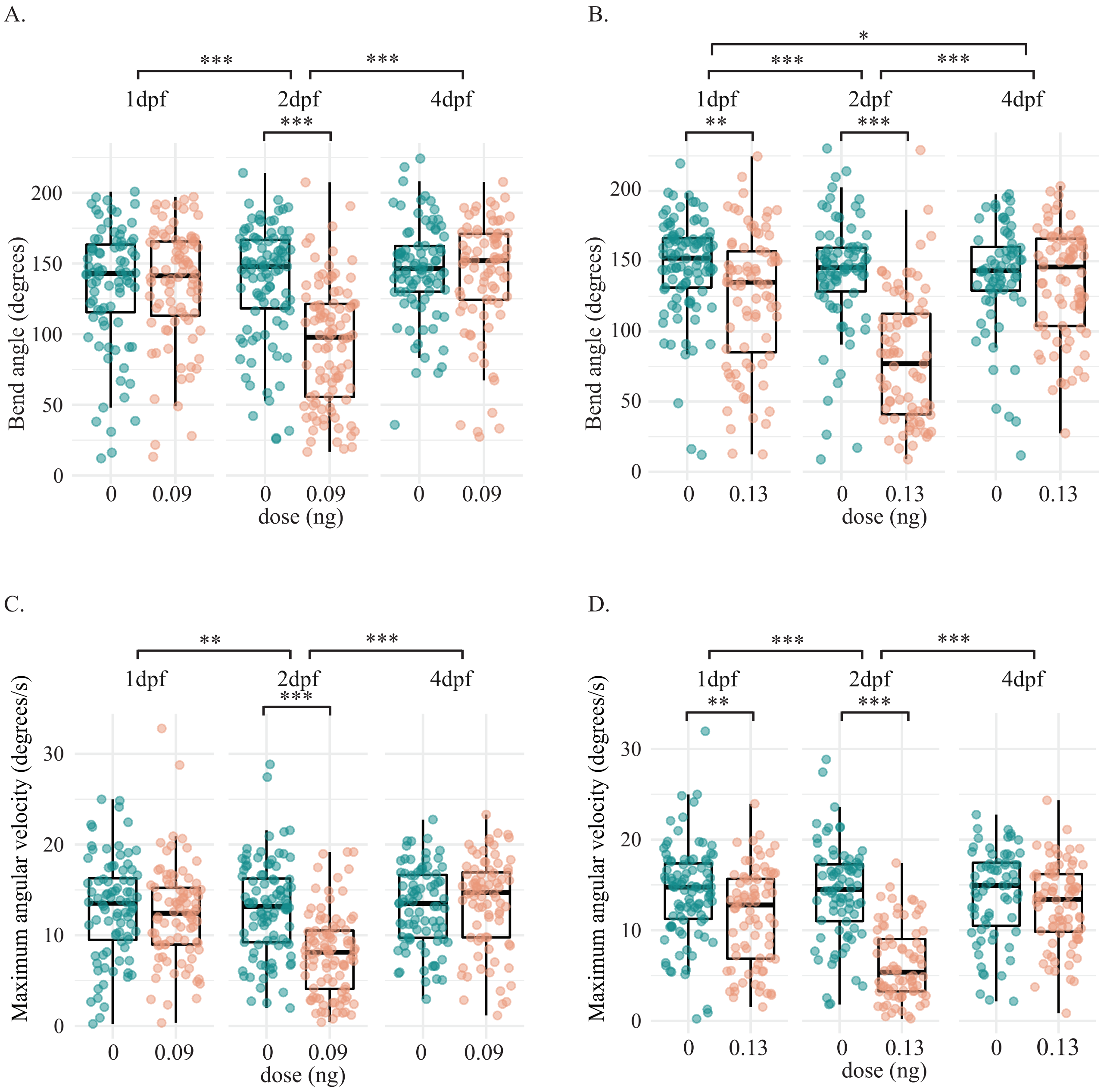


**Supplemental Figure 3. Exposure to domoic acid at 2 dpf (but not 1 or 4 dpf) alters LLC kinematics at the lowest dose.**
**(A)** Bend angles during LLC startles in fish exposed during different exposure days (1, 2 and 4 dpf) to the nominal dose of 0.09 ng of DomA and **(B)** to a dose that ranged from 0.126- 0.144 (labelled as 0.13ng of DomA). **(C)** Maximal angular velocity during LLC startles in fish exposed during different exposure days to 0.09 ng of DomA and to **(D)** to a dose that ranged from 0.126- 0.144 (labelled as 0.13ng of DomA). Each point represents the median kinematic response of an individual larvae to multiple identical stimuli.
* p <0.05, **=p<0.01, *** p< 0.001.

Figure supplement: Supplemental video 1 is a sample startle response.

| Sample confocal microscopy images | Sample widefield microscopy images | Severity Classification (number) - Description |
|---|---|---|
| 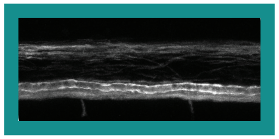 | 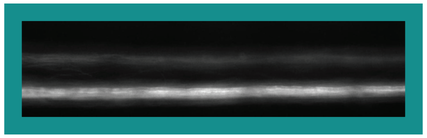 | (0) Normal phenotype — Dorsal and ventral regions of the spinal cord had labeled myelin sheaths. The myelin sheath surrounding the Mauthner axon was visible. |
| 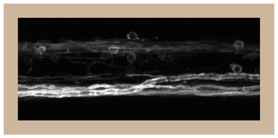 | 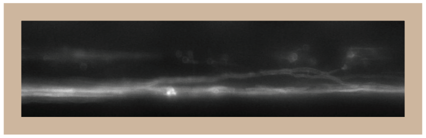 | (1) Myelin sheaths were present but disorganized. In some cases, myelinated axons that were normally found ventrally were located more dorsally. In others, the myelinated axons terminated prematurely with distal ends located more dorsally. |
| 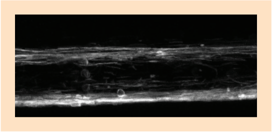 | 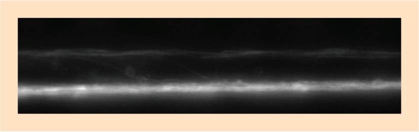 | (2) Myelin was labeled in both the dorsal and ventral regions of the spinal cord, but there are some noticeable deficits. While the ventral spinal cord was labeled, it had noticeably less myelin labeled compared to controls. |
| 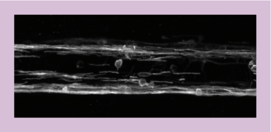 | 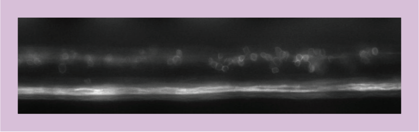 | (3) The loss of labeled myelin in the ventral spinal cord resulted in large, observable gaps between myelinated axons. |
| 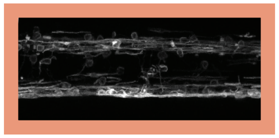 | 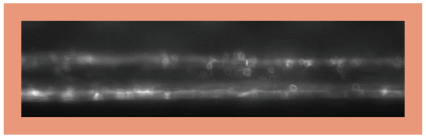 | (4) Myelin sheaths were essentially nonexistent in the ventral spinal cord. Instead, numerous hollow circular profiles were present both in the ventral and dorsal spinal cord. |
| | 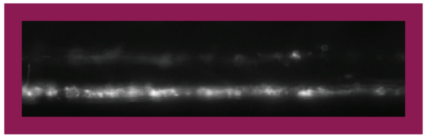 | (5) There were visible sloughed off portions of myelin which were defined by labeled portions of myelin which had a 'rough looking appearance' and were separated from the thin, elongated sheaths |

**Supplemental figure 4: Qualitative myelin phenotypes**
Myelinated axonal tracts in the dorsal and ventral spinal cord. Fish were blindly classified into 6 categories (0-5) based on the severity in the myelin defect. Representative confocal and widefield fluorescence microscopy images are shown for each severity classification.

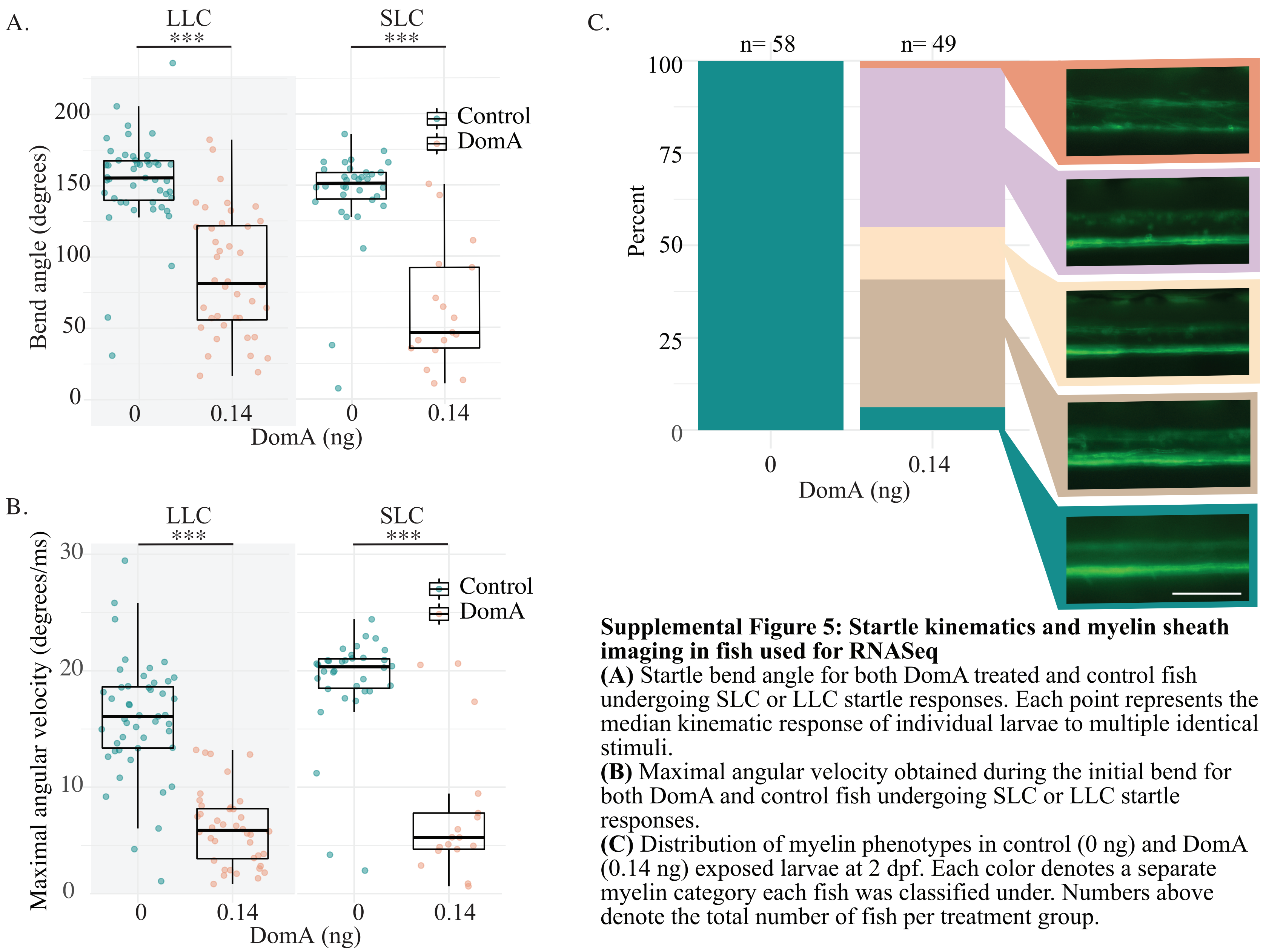


**Supplemental Figure 5: Startle kinematics and myelin sheath imaging in fish used for RNASeq**
**(A)** Startle bend angle for both DomA treated and control fish undergoing SLC or LLC startle responses. Each point represents the median kinematic response of individual larvae to multiple identical stimuli.
**(B)** Maximal angular velocity obtained during the initial bend for both DomA and control fish undergoing SLC or LLC startle responses.
**(C)** Distribution of myelin phenotypes in control (0 ng) and DomA (0.14 ng) exposed larvae at 2 dpf. Each color denotes a separate myelin category each fish was classified under. Numbers above denote the total number of fish per treatment group.

*** = $p < 0.0001$; Scale bar = 100 µm.

**SUPPLEMENTAL TABLES**

**Supplemental Table 1**: **Developmental time ranges in hours post fertilization (hpf) for fish included in each injection category**

| Injection label | Time range (hpf) |
|---|---|
| 1 dpf | 28-32.5 |
| 2 dpf | 47-52 |
| 4 dpf | 100- 105 |
| 1.5 dpf | 37-39 |
| 2.25 dpf | 54-55 |
| 2.50 dpf | 59.5- 60 |
| 2.5 dpf | 62-64 |
| 2.75 dpf | 66- 67 |
| 3 dpf | 71- 74.5 |

**Supplemental Table 2: Mortality in *Tg(mbp:EGFP-CAAX)* larvae exposed to DomA by intravenous injection**

| Day injected | Control | DomA (nominal dose) | | | |
|---|---|---|---|---|---|
| | **0 ng** | **0.09 ng** | **0.13 ng** | **0.14 ng** | **0.18 ng** |
| **1 dpf** | 0.4% ± 0.4 (n= 6) | 1.9% ± 1.3 (n= 3) | 3.1% | | 0.8% ± 0.5 (n= 5) |
| **1.5 dpf** | 0% (n= 1) | | | | 0% (n= 1) |
| **2 dpf** | 1.2% ± 0.7 (n= 13) | 0.4% ± 0.4 (n= 3) | 2.9%, 0% | 0% ± 0 (n= 6) | 6.2% ± 3.9 (n= 6) |
| **2.5 dpf** | 0% (n= 1) | | | 0% (n= 1) | |
| **3 dpf** | 0% (n= 1) | | | 0% (n= 1) | |
| **4 dpf** | 0.2% ± 0.2 (n= 8) | 0% (n= 3) | 0%, 0% | 0%, 0% | 0.9% ± 0.6 (n= 4) |

*Notes:* Mortality was recorded at 5 days post fertilization (dpf). n= number of repeated trials ± standard error (SE) of the mean when there are more than two repeated experiments. Otherwise, the raw percentage is listed per trial

**Supplemental Table 3: Trials included for the swim bladder analysis**

| | | Control | DomA (nominal dose) | | | |
|---|---|---|---|---|---|---|
| | | **0 ng** | **0.09 ng** | **0.13 ng** | **0.14 ng** | **0.18 ng** |
| **Day injected** | **1 dpf** | n= 20 | - | - | n= 24 | - |
| | **2 dpf** | n= 4, 25, 8, 26, 24, 47 | - | n= 9 | n= 38, 12, 7, 26, 27, 40 | n= 10 |
| | **4 dpf** | n= 25, 10 | n= 19, 9 | - | n= 25 | n= 12, 10 |

*Notes:* n corresponds to the number of fish within a trial that were exposed to a given dose of DomA administered at the specified developmental time period. Fish were imaged from 5-7 dpf. Fish injected at 4 dpf with 0.18 ng DomA that also had 'opaque brains' were excluded from this analysis (See Supplemental Table 4 for opaque brain phenotype breakdown).

**Supplemental Table 4: Opaque brains in *Tg(mbp:EGFP-CAAX)* larvae exposed to DomA by intravenous injection**

| Day injected | Control | DomA (nominal dose injected) | | | |
|---|---|---|---|---|---|
| | **0 ng** | **0.09 ng** | **0.13** | **0.14** | **0.18 ng** |
| 1 dpf | 0% ± 0 (n= 6) | 0% ± 0 (n= 3) | 0% (n= 1) | | 0% (n= 5) |
| 1.5dpf | (0%, 0%) | | | | 0% (n= 1) |
| 2 dpf | 0% ± 0 (n= 13) | 0% ± 0 (n= 3) | 0%, 0% | 0% ± 0 (n= 6) | 0% ± 0 (n= 6) |
| 2.5dpf | (0%, 0%) | | | 0% (n= 1) | |
| 3 dpf | 0% (n= 1) | | | 0% (n= 1) | |
| 4 dpf | 0% ± 0 (n= 8) | 0.5% ± 0.5 (n= 3) | 0%, 2.2% | (0%, 5%) | 40.6% ± 9.5 (n= 4) |

*Notes:* Percent of the population with opaque brains were recorded at 5 days post fertilization (dpf)
± = the standard error of the mean. n = number of trials included.

**Supplemental Table 5: Acute neurological phenotypes following developmental exposures to DomA at 1 dpf**

| Dose (ng) | Trials | Day observed | Convulsions mean % ± SE or (mean %, mean%) | No touch response mean % ± SE or (mean %, mean%) |
|---|---|---|---|---|
| 0 | n= 5 | 2 dpf | 0 | 0.4 ± 0.4 |
| | | 3 dpf | 0 | 0.2 ± 0.2 |
| | | 4 dpf | 0 | 0 |
| | | 5 dpf | 0 | 0 |
| 0.09 | n= 2 | 2 dpf | (2.9,0) | (32.4, 15.9) |
| | | 3 dpf | (0, 0) | (0,0) |
| | | 4 dpf | (0, 0) | (0,0) |
| | | 5 dpf | (0, 0) | (0,0) |
| 0.18 | n= 4 | 2 dpf | 8.0 ± 3.0 | 53.4 ± 12.9 |
| | | 3 dpf | 2.0 ± 1.6 | 17.3 ± 9.7 |
| | | 4 dpf | 0.8 ± 0.8 | 0.4 ± 0.4 |
| | | 5 dpf | 0 | 0 |

*Notes:* Percent of fish exhibiting convulsions or no touch responses tracked daily post exposure until 5 dpf (2-5 dpf). n = number of repeated trials ± SE of the mean when there are greater than two repeated experiments. Otherwise, the raw percentage is listed per trial. Within an individual treatment group, the minimum number of fish was 31 and the maximum number was 92.

**Supplemental Table 6: Acute neurological phenotypes following developmental exposures to DomA at 2 dpf**

| Dose (ng) | Trials | Day observed | Convulsion mean ± SE or (mean, mean) | No touch response mean ± SE or (mean, mean) |
|---|---|---|---|---|
| | | 3 dpf | 0 | 0.2 ± 0.2 |
| 0 | n= 11 | 4 dpf | 0 | 0.4 ± 0.3 |
| | | 5dpf | 0 | 0.1 ± 0.1 |
| | | 3 dpf | (2.4, 0) | (16.9, 11.7) |
| 0.09 | n = 2 | 4 dpf | (0,0) | (0, 1.3) |
| | | 5dpf | (0,0) | (0,0) |
| | | 3 dpf | 23.7 ± 11.6 | 38.8 ± 10.3 |
| 0.14 | n= 6 | 4 dpf | 0 | 1.5 ± 1.5 |
| | | 5dpf | 0 | 0.6 ± 0.6 |
| | | 3 dpf | 40.0 ± 7.4 | 47.8 ± 16.0 |
| 0.18 | n= 4 | 4 dpf | 0.4 ± 0.4 | 5.6 ± 3.6 |
| | | 5dpf | 1.4 ± 1.4 | 0.4 ± 0.4 |

*Notes:* Percent of fish exhibiting convulsions or no touch responses tracked over the course of development (3-5 dpf). n = number of repeated trials. ± SE of the mean when there are more than 2 repeated experiments. Otherwise, the raw percentage data is listed per trial. Within an individual treatment group, the minimum number of fish was 20 and the maximum number was 96.

**Supplemental Table 7: Acute neurological phenotypes following developmental exposures to DomA at 4 dpf**

| Dose (ng) | Trials | Convulsions mean % ± SE or (mean %, mean%) | No touch response mean % ± SE or (mean %, mean%) |
|---|---|---|---|
| 0 ng | n= 8 | 0 ± 0 | 0 ± 0 |
| 0.09 ng | n= 3 | 0 ± 0 | 0 ± 0 |
| 0.14 ng | n= 2 | (0, 0) | (0, 5.0) |
| 0.18 ng | n= 4 | 1.9 ± 1.9 | 24.1 ± 15.3 |

*Notes:* Percent of fish exhibiting convulsions or no touch responses assessed at 5 dpf. n = number of repeated trials ± SE of the mean when there are greater than two repeated experiments. Otherwise, the raw percentage is listed per trial. Within an individual treatment group, the minimum number of fish was 27 and the maximum number was 96.

**Supplemental Table 8: Model coefficients for the prevalence of the lack of touch responses for fish exposed at 1 dpf and 2 dpf**

| | Exposure at 1 dpf | Exposure at 2 dpf |
|---|---|---|
| | **Est Coef (Std. error)** | **Est Coef (Std. error)** |
| **Intercept** | -3.8 (0.5)*** | -4.2 (0.4) *** |
| **Dose** | +22.1 (3.9)*** | +23.7 (4.2)*** |
| **2 days post exposure** | -2.2 (0.7)*** | -2.9 (0.6)*** |
| **3 days post exposure** | -5.8 (1.3)** | -4.9 (0.6)*** |
| **4 days post exposure** | -46.3 (0.5)*** | *NA* |

*Notes:* Est Coef= estimated coefficient, Std. error = standard error ** = $p < 0.01$ *** = $p < 0.0001$

**Supplemental Table 9: Model coefficients for the prevalence of the convulsions and pectoral fin flapping for fish exposed at 1 dpf and 2 dpf**

| | Exposure at 1 dpf | Exposure at 2 dpf |
|---|---|---|
| | **Est Coef (Std. error)** | **Est Coef (Std. error)** |
| **Intercept** | -7.3 (1.2) *** | -6.6 (1.1) *** |
| **Dose** | +26.3 (6.8)*** | +35.1 (6.6)*** |
| **2 days post exposure** | -1.2 (0.8) | -5.4 (0.9)*** |
| **3 days post exposure** | -3.0 (0.9)** | -4.3 (0.7)*** |
| **4 days post exposure** | -44.8 (0.8)*** | *NA* |

*Notes:* Est Coef= estimated coefficient, Std. error = standard error, *** = $p < 0.0001$, *0.05 < $p$ < 0.01

**Supplemental Table 10: Post-hoc pairwise Dunnett comparisons following binomial modeling of percent responsiveness in startle behavior**

| | Exposures at 1 dpf | | Exposures at 2 dpf | | Exposures at 4 dpf | |
|---|---|---|---|---|---|---|
| **Dose comparisons (DomA - Control)** | **Coef (Std. Er)** | **Pr(>\|z\|)** | **Coef (Std. Er)** | **Pr(>\|z\|)** | **Coef (Std. Er)** | **Pr(>\|z\|)** |
| **0.18 ng – 0 ng == 0** | -0.8 (0.1) | **< 1e-4** | -0.7 (0.2) | **< 1e-8** | -0.9 (0.1) | **< 1e- 4** |
| **0.14 ng – 0 ng == 0** | -0.5 (0.1) | **0.001** | -0.8 (0.2) | **< 1e-8** | -0.2 (0.1) | 0.75 |
| **0.13 ng – 0 ng == 0** | -0.8 (0.2) | **< 1e-4** | -1.0 (0.2) | **< 1e-8** | 0.1 (0.2) | 0.98 |
| **0.09 ng – 0 ng == 0** | -0.1 (0.1) | 0.988 | -0.8 (0.2) | **< 1e-8** | 0.0 (0.1) | 0.997 |

*Notes*: Coef = estimated coefficient, Std. Er = standard error, Pr(>|z|) = p-value of the z-statistic. Significant values are in red font.

**Supplemental Table 11: Nonparametric analysis of SLC kinematics following exposure to different doses of DomA at 2 dpf**

| | | | Maximal Angular Velocity (SLC) | | | | Bend angle (SLC) | | | |
|---|---|---|---|---|---|---|---|---|---|---|
| **dose (ng)** | **trial day** | **n (SLC)** | **Est** | **95% CI [Lower, Upper]** | **Stat** | **p** | **Est** | **95% CI [Lower, Upper]** | **Stat** | **p** |
| **0.09** | 5/30/16 | 0 ng (n= 24), 0.13ng (n= 40) | 0.209 | [0.073, 0.346] | -4.877 | **2.98e-5** | 0.20 | [0.068, 0.332] | -5.18 | **1.48e-5** |
| | 5/19/16 | 0 ng (n=37), 0.09ng (n= 36) | 0.115 | [0.015, 0.214] | -8.987 | **2.56e-10** | 0.116 | [0.022, 0.211] | -9.1 | **1.15e-12** |
| **0.13** | 5/30/16 | 0 ng (n= 24), 0.13ng (n= 35) | 0.107 | [-0.007, 0.221] | -7.877 | **1.41e-9** | 0.11 | [0.004, 0.215] | -8.41 | **6.65e-10** |
| | 6/24/16 | 0 ng (n= 21), 0.13ng (n= 9) | 0.001 | [0, 0.002] | -1010 | **< 1e-14** | 0.001 | [0, 0.002] | -1010 | **< 1e-14** |
| **0.14** | 6/24/16 | 0 ng (n= 21), 0.14 ng (n =11) | 0.001 | [0, 0.002] | -1010 | **< 1e-14** | 0.001 | [0, 0.002] | -1010 | **< 1e-14** |
| | 10/10/16 | 0 ng (n= 34), 0.14 ng (n= 17) | 0.112 | [0.004, 0.221] | -7.252 | **< 1e-14** | 0.092 | [0, 0.184] | -8.94 | **< 1e-14** |
| | 9/23/16 | 0 ng (n= 42), 0.14 ng (n= 31) | 0.06 | [-0.006, 0.126] | 13.33 | **< 1e-14** | 0.029 | [-0.008, 0.067] | -25.08 | **< 1e-14** |
| **0.18** | 5/19/16 | 0 ng (n=37), 0.18ng (n= 26) | 0.169 | [0.023, 0.316] | -5.253 | **1.49e-5** | 0.09 | [0.002, 0.179] | -10.40 | **8.10e-15** |
| | 4/27/16 | 0 ng (n=17), 0.18ng (n= 10) | 0.071 | [-0.056, 0.197] | -7.183 | **< 1e-14** | 0.094 | [-0.039, 0.228] | -6.32 | **< 1e-14** |
| | 3/27/16 | 0 ng (n= 23), 0.18ng (n=19) | 0.005 | [-0.007,0.016] | -89.063 | **< 1e-14** | 0.002 | [-0.004, 0.009] | -153.80 | **< 1e-14** |
| | 4/15/16 | 0 ng (n=8), 0.18ng (n= 8) | 0.001 | [-0.172, 0.174] | -5.646 | **< 1e-14** | 0.001 | [-0.172, 0.174] | -5.646 | **< 1e-14** |

*Notes:* Est = relative estimated effect, Stat = statistic, p = p-value. Significant values are in red font.

**Supplemental Table 12: Nonparametric analysis of LLC kinematics following exposure to different doses of DomA at 2 dpf**

| | | | Maximal Angular Velocity (LLC) | | | | Bend angle (LLC) | | | |
|---|---|---|---|---|---|---|---|---|---|---|
| **dose (ng)** | **trial day** | **n (LLC)** | **Est** | **95% CI [Lower, Upper]** | **Stat** | **p** | **Est** | **95% CI [Lower, Upper]** | **Stat** | **p** |
| **0.09** | 5/30/16 | 0 ng (n= 40), 0.09 ng (n= 48) | 0.21 | [0.099, 0.321] | -5.892 | **1.53e-7** | 0.238 | [0.121, 0.354] | -5.078 | **4.51e-6** |
| | 5/19/16 | 0 ng (n=56), 0.09 ng (n= 47) | 0.253 | [0.143, 0.363] | -5.027 | **4.64e-6** | 0.207 | [0.105, 0.309] | -6.425 | **1.18e-8** |
| **0.13** | 5/30/16 | 0 ng (n= 40), 0.13 ng (n= 48) | 0.13 | [0.046, 0.215] | -9.904 | **1.33e-15** | 0.161 | [0.062, 0.260] | -7.734 | **4.19e-11** |
| | 6/24/16 | 0 ng (n= 24), 0.13 ng (n= 30) | 0.044 | [-0.025, 0.114] | -14.615 | **2.24e-14** | 0.062 | [-0.019, 0.144] | -12.182 | **1.98e-12** |
| **0.14** | 6/24/16 | 0 ng (n= 24), 0.14 ng (n = 30) | 0.043 | [-0.028,0.114] | -14.24 | **4.22e-14** | 0.054 | [-0.021, 0.129] | -13.41 | **1.99e-13** |
| | 10/10/16 | 0 ng (n= 47), 0.14 ng (n= 40) | 0.062 | [0.005, 0.118] | -15.618 | **<1e-14** | 0.107 | [0.029, 0.185] | -10.078 | **<1e-14** |
| | 9/23/16 | 0 ng (n= 71), 0.14 ng (n= 65) | 0.073 | [0.028, 0.119] | -18.491 | **<1e-14** | 0.085 | [0.038,0.132] | -17.458 | **<1e-14** |
| **0.18** | 5/19/16 | 0 ng (n=56), 0.18 ng (n=58) | 0.233 | [0.127, 0.340] | -5.61 | **3.96e-7** | 0.201 | [0.104, 0.298] | -6.89 | **1.41e-9** |
| | 4/27/16 | 0 ng (n= 19), 0.18 ng (n=20) | 0.108 | [-0.014, 0.23] | -6.61 | **<1e-14** | 0.142 | [-0.004, 0.288] | -4.976 | **<1e-14** |
| | 3/27/16 | 0 ng (n= 28), 0.18 ng (n = 26) | 0.074 | [-0.01, 0.159] | -10.298 | **<1e-14** | 0.11 | [0.009, 0.211] | -7.725 | **<1e-14** |
| | 4/15/16 | 0 ng (n=18), 0.18 ng (n= 33) | 0.059 | [-0.045, 0.162] | -8.966 | **<1e-14** | 0.072 | [-0.042, 0.187] | -7.835 | **<1e-14** |

*Notes*: Est = relative estimated effect, Stat = statistic, p = p-value. Significant values are in red font.

**Supplemental Table 13: Median and Interquartile range for startle kinematic parameters of fish exposed to different doses of DomA at 2 dpf**

| | | LLC | | | | SLC | | | |
|---|---|---|---|---|---|---|---|---|---|
| | | Bend angle | | Maximal angular velocity | | Bend angle | | Maximal angular velocity | |
| Dose (ng) | Trial day | Median | IQR | Median | IQR | Median | IQR | Median | IQR |
| 0 | 03_27_2016 | 142.74 | 28.39 | 15.36 | 3.55 | 139.48 | 12.51 | 20.07 | 1.93 |
| | 04_15_2016 | 134.79 | 44.27 | 11.70 | 4.36 | 147.88 | 15.51 | 19.93 | 3.71 |
| | 04_27_2016 | 153.34 | 18.79 | 15.97 | 5.05 | 150.72 | 17.73 | 19.03 | 2.83 |
| | 05_19_2016 | 156.25 | 63.49 | 13.27 | 8.64 | 156.78 | 24.74 | 19.50 | 1.80 |
| | 05_30_2016 | 143.56 | 29.35 | 13.17 | 6.22 | 148.01 | 26.58 | 19.79 | 1.83 |
| | 06_24_2016 | 180.46 | 30.42 | 17.14 | 5.14 | 163.25 | 15.73 | 19.72 | 1.55 |
| | 09_04_2016 | 151.4 | 29.91 | 16.44 | 4.74 | 146.47 | 19.81 | 18.48 | 3.05 |
| | 09_23_2016 | 149.24 | 35.88 | 15.32 | 4.73 | 150.68 | 21.75 | 20.08 | 1.96 |
| | 10_10_2016 | 155.43 | 27.64 | 16.08 | 5.26 | 151.30 | 18.59 | 20.33 | 2.53 |
| 0.09 | 05_19_2016 | 97.4 | 61.15 | 8.85 | 5.99 | 90.33 | 60.19 | 11.51 | 7.27 |
| | 05_30_2016 | 101.51 | 76.40 | 7.24 | 5.78 | 102.85 | 78.20 | 10.59 | 12.63 |
| 0.13 | 05_30_2016 | 76.57 | 74.10 | 4.94 | 5.90 | 74.69 | 51.29 | 10.33 | 9.42 |
| | 06_24_2016 | 83.02 | 55.10 | 5.67 | 3.01 | 49.15 | 36.17 | 5.09 | 2.13 |
| 0.14 | 06_24_2016 | 95.69 | 46.50 | 6.13 | 3.47 | 64.37 | 29.90 | 6.27 | 2.87 |
| | 09_23_2016 | 78.58 | 45.61 | 5.8 | 4.01 | 58.06 | 44.90 | 5.73 | 3.53 |
| | 10_10_2016 | 81.29 | 66.03 | 6.3 | 4.29 | 46.50 | 56.58 | 5.69 | 3.11 |
| 0.18 | 03_27_2016 | 75.02 | 63.27 | 5.68 | 4.55 | 56.61 | 61.86 | 6.66 | 7.27 |
| | 04_15_2016 | 59.56 | 39.59 | 4.26 | 2.41 | 37.12 | 17.42 | 4.59 | 2.90 |
| | 04_27_2016 | 49.77 | 59.17 | 4.00 | 2.52 | 54.28 | 42.04 | 6.98 | 4.98 |
| | 05_19_2016 | 71.3 | 58.68 | 6.60 | 3.97 | 67.48 | 79.20 | 6.38 | 10.59 |

*Note:* IQR = interquartile range, 3rd quantile- 1st quantile

**Supplemental Table 14: Nonparametric analysis of SLC kinematics following exposure to different doses of DomA at 1 dpf**

| | | | Maximal Angular Velocity (SLC) | | | | Bend angle (SLC) | | | |
|---|---|---|---|---|---|---|---|---|---|---|
| **dose (ng)** | **trial day** | **n (SLC)** | **Est** | **95% CI [Lower, Upper]** | **Stat** | **p** | **Est** | **95% CI [Lower, Upper]** | **Stat** | **p** |
| **0.09** | 5/30/16 | 0 ng (n= 39), 0.09 ng (n= 36) | 0.555 | [0.391, 0.719] | 0.800 | 0.669 | 0.426 | [0.268, 0.584] | -1.10 | 0.469 |
| | 5/19/16 | 0 ng (n=25), 0.09 ng (n= 25) | 0.419 | [0.226, 0.613] | -0.96 | 0.532 | 0.581 | [0.387, 0.774] | 0.96 | 0.536 |
| **0.13** | 5/30/16 | 0 ng (n= 39), 0.13 ng (n=19) | 0.356 | [0.140, 0.573] | -1.59 | 0.232 | 0.309 | [0.121, 0.497] | -2.39 | **0.045** |
| **0.14** | 9/4/16 | 0 ng (n=35), 0.14 ng(n=35) | 0.557 | [0.416, 0.697] | 0.81 | 0.423 | 0.347 | [0.21,0.484] | -2.24 | **0.029** |
| **0.18** | 5/19/16 | 0 ng (n=25), 0.18 ng (n=18) | 0.449 | [0.235,0.663] | -0.55 | 0.805 | 0.549 | [0.336, 0.762] | 0.53 | 0.82 |
| | 4/27/16 | 0 ng (n=13), 0.18 ng (n= 16) | 0.279 | [0.078, 0.48] | -2.26 | **0.032** | 0.274 | [0.077,0.471] | -2.36 | **0.026** |
| | 4/15/16 | 0 ng (n=12), 0.18 ng (n= 12) | 0.34 | [0.088, 0.593] | -1.32 | 0.203 | 0.222 | [0.01, 0.435] | -2.71 | **0.013** |

*Notes:* Trials with multiple doses were analyzed using Dunnett-type intervals to compare kinematics of fish exposed at each dose to the controls. Trials with single doses were tested using nonparametric t-tests. Est = relative estimated effect, Stat = statistic, p = p-value. Significant values are in red font.

**Supplemental Table 15: Nonparametric analysis of LLC kinematics following exposure to different doses of DomA at 1 dpf**

| | | | Maximal Angular Velocity (LLC) | | | | Bend angle (LLC) | | | |
|---|---|---|---|---|---|---|---|---|---|---|
| **dose (ng)** | **trial day** | **n (LLC)** | **Est** | **95% CI [Lower, Upper]** | **Stat** | **p** | **Est** | **95% CI [Lower, Upper]** | **Stat** | **p** |
| **0.09** | 5/30/16 | 0 ng (n= 47), 0.09 ng (n= 51) | 0.429 | [0.296, 0.562] | -1.212 | 0.39 | 0.476 | [0.340, 0.613] | -0.398 | 0.901 |
| | 5/19/16 | 0 ng (n= 37), 0.09 ng (n= 35) | 0.492 | [0.334, 0.650] | -0.115 | 0.99 | 0.544 | [0.388, 0.699] | 0.6313 | 0.742 |
| **0.13** | 5/30/16 | 0 ng (n= 47), 0.13 ng (n= 31) | 0.298 | [0.156, 0.440] | -3.24 | **0.004** | 0.321 | [0.163, 0.478] | -2.63 | **0.023** |
| **0.14** | 9/4/16 | 0 ng (n=50), 0.14 ng (n= 48) | 0.385 | [0.271, 0.498] | -2.02 | **0.046** | 0.335 | [0.224, 0.447] | -2.941 | **0.004** |
| **0.18** | 5/19/16 | 0 ng (n= 37), 0.18 ng (n= 41) | 0.372 | [0.228, 0.517] | -1.984 | 0.09 | 0.405 | [0.258, 0.552] | -1.454 | 0.249 |
| | 4/27/16 | 0 ng (n= 15), 0.18 ng (n= 17) | 0.118 | [-0.012, 0.247] | -6.087 | **< 1e-14** | 0.11 | [-0.004, 0.224] | -6.997 | **< 1e-14** |
| | 4/15/16 | 0 ng (n= 36), 0.18 ng (n= 37) | 0.354 | [0.221, 0.486] | -2.214 | **0.031** | 0.322 | [0.196, 0.448] | -2.817 | **0.006** |

*Notes*: Trials with multiple doses were analyzed using Dunnett-type intervals to compare the kinematics of fish exposed at each dose to the control. Trials with single DomA doses were tested using nonparametric t-tests. Est = relative estimated effect, Stat = statistic, p = p-value. Significant values are in red font.

**Supplemental Table 16: Nonparametric analysis of SLC kinematics following exposure to different doses of DomA at 4 dpf.**

| | | | Maximal Angular Velocity (SLC) | | | | Bend angle (SLC) | | | |
|---|---|---|---|---|---|---|---|---|---|---|
| **dose (ng)** | **trial day** | **n (SLC)** | **Est** | **95% CI [Lower, Upper]** | **Stat** | **p** | **Est** | **95% CI [Lower, Upper]** | **Stat** | **p** |
| 0.09 | 5/30/16 | 0 ng (n= 31), 0.09 ng (n= 48) | 0.544 | [0.390, 0.697] | 0.642 | 0.741 | 0.545 | [0.394, 0.696] | 0.676 | 0.730 |
| | 5/19/16 | 0 ng (n=20), 0.09 ng (n = 17) | 0.435 | [0.181, 0.690] | -0.624 | 0.781 | 0.435 | [0.196, 0.675] | -0.654 | 0.756 |
| 0.13 | 5/30/16 | 0 ng (n= 31), 0.09 ng (n= 45) | 0.466 | [0.315, 0.617] | -0.509 | 0.826 | 0.448 | [0.296, 0.600] | -0.776 | 0.662 |
| | 6/24/16 | 0 ng (n= 5), 0.13 ng (n = 11) | 0.618 | [0.240, 0.996] | 0.778 | 0.68 | 0.600 | [0.214, 0.986] | 0.651 | 0.750 |
| 0.14 | 6/24/16 | 0 ng (n= 5), 0.14 ng(n= 13) | 0.585 | [0.239, 0.930] | 0.61 | 0.792 | 0.338 | [-0.009,-0.686] | -1.167 | 0.429 |
| 0.18 | 5/19/16 | 0 ng (n=20), 0.18 ng (n= 10) | 0.430 | [0.138, 0.722] | -0.588 | 0.802 | 0.37 | [0.097, 0.643] | -1.15 | 0.441 |
| | 4/27/16 | 0 ng (n= 13), 0.18 ng (n= 8) | 0.212 | [-0.058, 0.481] | -2.309 | **0.038** | 0.183 | [-0.026, 0.391] | -3.185 | **0.005** |
| | 3/27/16 | 0 ng (n=19), 0.18 ng (n= 10) | 0.363 | [0.107, 0.62] | -1.143 | 0.272 | 0.389 | [0.15, 0.629] | -0.967 | 0.346 |

*Note:* Significant values are in red font.

**Supplemental Table 17: Nonparametric analysis of LLC kinematics following exposure to different doses of DomA at 4 dpf**

| | | | Maximal Angular Velocity (LLC) | | | | Bend angle (LLC) | | | |
|---|---|---|---|---|---|---|---|---|---|---|
| **dose (ng)** | **trial day** | **n (LLC)** | **Est** | **95% CI [Lower, Upper]** | **Stat** | **p** | **Est** | **95% CI [Lower, Upper]** | **Stat** | **p** |
| 0.09 | 5/30/16 | 0 ng (n= 37), 0.09 ng (n= 45) | 0.499 | [0.353, 0.645] | -0.014 | 1.000 | 0.546 | [0.399, 0.693] | 0.706 | 0.713 |
| | 5/19/16 | 0 ng (n= 47), 0.09 ng (n= 36) | 0.592 | [0.442, 0.741] | 1.441 | 0.289 | 0.472 | [0.321, 0.623] | -0.430 | 0.887 |
| 0.13 | 5/30/16 | 0 ng (n= 37), 0.13 ng (n= 47) | 0.511 | [0.365, 0.657] | 0.173 | 0.978 | 0.56 | [0.413, 0.706] | 0.921 | 0.568 |
| | 6/24/16 | 0 ng (n=18), 0.13 ng (n = 33) | 0.283 | [0.102, 0.464] | -2.763 | **0.017** | 0.226 | [0.059, 0.392] | -3.809 | **0.001** |
| 0.14 | 6/24/16 | 0 ng (n=18), 0.14 ng(n= 22) | 0.328 | [0.123, 0.533] | -1.93 | 0.1089 | 0.283 | [0.085, 0.481] | -2.529 | **0.030** |
| 0.18 | 5/19/16 | 0 ng (n= 47), 0.18 ng (n= 26) | 0.394 | [0.203, 0.585] | -1.309 | 0.355 | 0.337 | [0.163, 0.511] | -2.184 | 0.069 |
| | 4/27/16 | 0 ng (n= 11), 0.18 ng (n= 11) | 0.504 | [0.215, 0.793] | 0.03 | 0.976 | 0.413 | [0.144, 0.682] | -0.676 | 0.507 |
| | 3/27/16 | 0 ng (n=14), 0.18 ng (n= 12) | 0.482 | [0.238,0.726] | -0.151 | 0.881 | 0.411 | [0.171, 0.65] | -0.769 | 0.449 |

*Note:* Significant values are in red font.

**Supplemental Table 18: Trials included to assess myelin labeling, imaged using confocal microscopy at 5 dpf**

| | Control | DomA |
|---|---|---|
| **1 dpf** | n= 15, 7 | n= 25, 6 |
| **1.5 dpf** | n= 18, 6 | n= 23,11 |
| **2 dpf** | n= 7, 8, 24, 22 | n= 31, 32, 36, 7 |
| **2.5 dpf** | n = 21, 8 | n= 33, 7 |
| **4 dpf** | n= 6,10, 24 | n= 14, 24, 8 |

*Notes:* n corresponds to the number of fish within a trial that were exposed to DomA administered at the specified developmental time period (leftmost column). DomA-exposed larvae were given a nominal dose that ranged from 0.13- 0.14 ng.

**Supplemental Table 19: Trials included to assess myelin labeling, imaged using widefield epifluorescence microscopy at 5 dpf**

| | | Control | DomA | | | |
|---|---|---|---|---|---|---|
| | | **0 ng** | **0.09 ng** | **0.13 ng** | **0.14 ng** | **0.18 ng** |
| **Day injected** | **1 dpf** | n=24, 12, 6, 18, 64, 37 | n=25, 16, 11 | n=6 | n= 68, 40 | n=28, 51,15 |
| | **1.5 dpf** | n= 30 | - | - | - | n= 48 |
| | **2 dpf** | n= 23, 37, 29, 19, 29, 70, 82, 80, 71, 30, 21, 80, 58 | n=51, 31, 13 | n=12, 34, 43 | n=35, 68, 75, 77, 81, 40, 24, 80, 49 | n= 26, 41, 59, 27 |
| | **2.5 dpf** | n= 18, 55, 17 | - | - | n= 20, 62, 36 | - |
| | **3 dpf** | n = 6, 52, 45 | - | n= 12 | n= 8,77,36 | - |
| | **4 dpf** | n= 14, 17, 18, 19, 50, 24 | n=17, 10 | n= 10, 32 | n= 19, 27, 48, 77, 21 | - |

*Note:* n corresponds to the number of fish within a trial that were exposed to a given dose of DomA administered at the specified developmental time period.

**Supplemental Table 20: Trials included to assess myelin labeling, imaged using widefield microscopy at 6 dpf**

| | Control | DomA | | |
|---|---|---|---|---|
| **Day injected** | **0 ng** | **0.09 ng** | **0.13 ng** | **0.18 ng** |
| **1 dpf** | n= 27, 10, 32 | n= 22, 36 | n= 36 | n= 25, 14 |
| **2 dpf** | n= 22, | n= 52 | n= 41 | - |
| **4 dpf** | n= 31, 24, 34 | n= 35, 13, 41 | n= 37 | - |

*Note:* n corresponds to the number of fish within a trial that were exposed to a given dose of DomA administered at the specified developmental time period.

**Supplemental Table 21: Trials included to assess myelin labeling, imaged using widefield microscopy at 7 dpf**

| | Control | DomA | | | |
|---|---|---|---|---|---|
| **Day injected** | **0 ng** | **0.09 ng** | **0.13 ng** | **0.14 ng** | **0.18 ng** |
| **1 dpf** | n=23 | n= 18 | n=17 | - | - |
| **2 dpf** | n=14, 10, 2, 28 | n=18 | n=19 | n=39 | n=19, 21 |
| **4 dpf** | n=21, 16 | n=19 | n=20 | n=25 | - |

*Note:* n corresponds to the number of fish within a trial that were exposed to a given dose of DomA administered at the specified developmental time period.

**Supplemental Table 22: Multinomial logistic regression model for distribution of myelin phenotypes in fish exposed to 0.14 ng of DomA at different periods in development**

| Myelin category | Intercept | 2 dpf inj | 2.5 dpf inj | 3 dpf inj | 4 dpf inj |
|---|---|---|---|---|---|
| 1 | -4.63 (p= 3.98e-6) | 6.50 (p= 3.76e-10) | 4.47 (p= 2.35e-5) | 1.00 (p= 4.19e-1) | -9.42 (p= 0.91) |
| 2 | -3.54 (p= 1.56e-9) | 5.02 (p= 6.66e-15) | 4.10 (p= 2.83e-10) | 2.04 (p= 1.55e-3) | -19.65 (p< e-16) |
| 3 | -31.83 (p= 7.00e-11) | 34.58 (p= 1.44e-12) | 32.54 (p= 2.71e-11) | 29.57 (p= 1.46e-9) | 20.19 (p= 0.30) |
| 4 | -4.64 (p= 4.00e-6) | 5.88 (p= 1.68e-8) | 2.74 (p= 2.04e-2) | 1.00 (p= 4.19e-1) | -7.74 (p= 0.83) |
| 5 | -31.01 (p< e-16) | 11.57 (p< e-16) | 28.70 (p< e-16) | 29.45 (p< e -16) | 26.45 (p< e-16) |

*Note:* The reference condition for the analysis is 1 dpf injected with no myelin deficits (myelin category 0). Coefficients were listed along with p-values in parentheses.

**Supplemental Table 23: Multinomial logistic regression model for distribution of myelin phenotypes in fish exposed to different doses of DomA at 2 dpf and imaged at 5, 6, and 7 dpf**

| myelin category | 5 dpf imaged | | 6 dpf imaged | | 7 dpf imaged | |
|---|---|---|---|---|---|---|
| | Intercept | DomA (ng) | Intercept | DomA (ng) | Intercept | DomA (ng) |
| 1 | -4.95 (p <e-16) | 54.07 (p <e-16) | -3.57 (p =0.0004) | 38.27 (p =0.0001) | -2.74 (p =1.61e-7) | 34.89 (p =6.07e-11) |
| 2 | -5.51 (p <e-16) | 54.36 (p <e-16) | -3.33 (p =0.0007) | 31.98 (p =0.001) | -4.26 (p =8.23e-7) | 42.14 (p =5.73e-9) |
| 3 | -5.04 (p <e-16) | 60.28 (p <e-16) | -6.84 (p =0.0003) | 61.59 (p =0.0003) | -5.12 (p =2.58e-7) | 49.97 (p =2.85e-10) |
| 4 | -7.84 (p <e-16) | 71.17 (p <e-16) | -5.26 (p =0.025) | 31.78 (p =0.159) | -15.92 (p =3.09e-2) | 100.40 (p =1.73e-2) |
| 5 | -8.86 (p =0.003) | 43.03 (p =0.07) | NA | NA | NA | NA |

*Notes:* DomA is modeled as a continuous factor from fish exposed to the following nominal DomA doses: 0.09, 0.126, 0.135, and 0.144 ng. Coefficients were listed along with p-values in parentheses.

**Supplemental Table 24: Genes associated with the enriched GO term: biological processes**

| GO term | term ID | pvalue | ENSEMBL_gene | gene name | logFC |
|---|---|---|---|---|---|
| Protein depolymerization | GO:0051261 | 3.49E-02 | ENSDARG00000030106 | *stmn4* | 0.507 |
| Microtubule depolymerization | GO:0007019 | 8.58E-03 | ENSDARG00000038465 | *stmn3* | -0.612 |
| | | | ENSDARG00000043932 | *stmn4l* | 0.433 |

**Supplemental Table 25: Human phenology phenotypes associated with differentially expressed genes at 3 dpf**

| **Human phenology phenotype** | **Term ID** | **pvalue** | ENSDARG00000012426 *neflb* | ENSDARG00000057568 *nefla* | ENSDARG00000038609 *mpz* | ENSDARG00000039522 *tubb2* | ENSDARG00000018997 *cplx2l* |
|---|---|---|---|---|---|---|---|
| Peripheral axonal degeneration | HP:0000764 | 0.02 | ■ | ■ | ■ | ■ | |
| Myelin outfoldings | HP:0004336 | 0.00 | ■ | ■ | ■ | | |
| Segmental peripheral demyelination/ remyelination | HP:0003481 | 0.01 | ■ | ■ | ■ | | |
| Clusters of axonal regeneration | HP:0007233 | 0.01 | ■ | ■ | | | |
| Ulnar claw | HP:0001178 | 0.00 | ■ | ■ | ■ | | |
| Hypotrophy of small hand muscles | HP:0006006 | 0.05 | ■ | ■ | ■ | | |
| Ectrodactyly | HP:0100257 | 0.04 | ■ | ■ | ■ | | ■ |
| Split hand | HP:0001171 | 0.04 | ■ | ■ | ■ | | ■ |

*Notes:* Genes associated with each human phenology phenotype are listed by ENSEMBL gene ID and common gene name. Shading indicates which of the listed genes are associated with each phenotype.

**Supplemental Table 26: Differentially expressed genes in DomA-exposed fish at 3 dpf (exposed at 2 dpf then tissue extracted at 3 dpf or 28 hours post exposure)**

| ENSEMBL_gene | Gene name | logFC | logCPM | F | PValue | FDR |
|---|---|---|---|---|---|---|
| ENSDARG00000031588 | *si:dkey-239b22.1* | 3.460 | 3.901 | 255.669 | 1.47e-10 | 3.08e-06 |
| ENSDARG00000077799 | *egr4* | -1.491 | 5.356 | 157.207 | 1.53e-09 | 1.59e-05 |
| ENSDARG00000028804 | *ankrd9* | -1.074 | 5.434 | 129.297 | 6.06e-09 | 4.22e-05 |
| ENSDARG00000074311 | *mast2* | 0.964 | 6.326 | 122.873 | 8.64e-09 | 4.51e-05 |
| ENSDARG00000097110 | *si:dkey-56f14.4* | -0.973 | 4.725 | 115.655 | 1.32e-08 | 5.49e-05 |
| ENSDARG00000027744 | *gadd45ba* | -0.921 | 7.133 | 97.579 | 4.21e-08 | 0.000146422 |
| ENSDARG00000012426 | *neflb* | -0.943 | 4.326 | 89.623 | 7.46e-08 | 0.000222491 |
| ENSDARG00000041140 | *ddb2* | -0.624 | 6.277 | 84.308 | 1.12e-07 | 0.000292604 |
| ENSDARG00000097528 | *si:dkey-7j14.5* | -0.727 | 6.256 | 82.711 | 1.27e-07 | 0.000295223 |
| ENSDARG00000043697 | *nefmb* | -1.076 | 5.258 | 83.299 | 1.74e-07 | 0.000362796 |
| ENSDARG00000021351 | *nefma* | -0.803 | 5.597 | 77.039 | 2.03e-07 | 0.000385537 |
| ENSDARG00000012281 | *zgc:65851* | -1.016 | 4.264 | 70.095 | 3.75e-07 | 0.000629667 |
| ENSDARG00000004187 | *zgc:122979* | -0.939 | 5.832 | 70.292 | 3.92e-07 | 0.000629667 |
| ENSDARG00000029795 | *fam213b* | -0.602 | 5.455 | 58.698 | 1.16e-06 | 0.001724981 |
| ENSDARG00000075891 | *sall1b* | 1.805 | 2.480 | 57.120 | 1.40e-06 | 0.00194405 |
| ENSDARG00000052917 | *si:ch211-202f3.3* | 0.719 | 4.502 | 52.467 | 2.31e-06 | 0.002836386 |
| ENSDARG00000078567 | *lonrf1l* | -0.675 | 6.601 | 52.458 | 2.31e-06 | 0.002836386 |
| ENSDARG00000074390 | *tmem176l.4* | 0.819 | 5.943 | 53.861 | 2.72e-06 | 0.002836386 |
| ENSDARG00000057568 | *nefla* | -1.326 | 3.659 | 52.984 | 2.76e-06 | 0.002836386 |
| ENSDARG00000102899 | *cremb* | 2.204 | 1.392 | 50.913 | 2.77e-06 | 0.002836386 |
| ENSDARG00000094860 | *gpr186* | -1.053 | 4.520 | 51.072 | 2.86e-06 | 0.002836386 |
| ENSDARG00000019498 | *cry5* | -0.718 | 5.029 | 50.294 | 2.99e-06 | 0.002836386 |
| ENSDARG00000087303 | *cebpd* | -0.594 | 8.725 | 49.658 | 3.23e-06 | 0.002929668 |
| ENSDARG00000053475 | *ngb* | -0.590 | 5.084 | 48.579 | 3.68e-06 | 0.003203961 |
| ENSDARG00000104919 | *si:ch211-153b23.3* | 2.442 | 1.688 | 48.068 | 4.90e-06 | 0.004090186 |
| ENSDARG00000037910 | *No longer in database* | 0.583 | 5.730 | 45.645 | 5.34e-06 | 0.004285939 |
| ENSDARG00000040135 | *fosaa* | -1.080 | 3.044 | 43.812 | 6.79e-06 | 0.005252425 |
| ENSDARG00000075048 | *lonrf1* | -0.510 | 7.065 | 44.833 | 9.23e-06 | 0.006667837 |
| ENSDARG00000103199 | *si:dkey-247k7.2* | 1.346 | 5.359 | 57.858 | 9.26e-06 | 0.006667837 |
| ENSDARG00000031702 | *prkg1b* | -0.890 | 5.910 | 45.425 | 9.66e-06 | 0.006726869 |
| ENSDARG00000030106 | *stmn4* | 0.507 | 6.716 | 40.426 | 1.08e-05 | 0.007291357 |
| ENSDARG00000089429 | *si:dkey-205h13.2* | -0.511 | 7.533 | 41.178 | 1.35e-05 | 0.00840476 |
| ENSDARG00000038639 | *elovl6l* | -0.725 | 5.040 | 38.875 | 1.35e-05 | 0.00840476 |
| ENSDARG00000074642 | *No longer in database* | 1.935 | 2.101 | 39.526 | 1.37e-05 | 0.00840476 |
| ENSDARG00000082789 | *NC_002333.18* | -0.655 | 4.798 | 37.108 | 1.76e-05 | 0.010498015 |
| ENSDARG00000091715 | *No longer in database* | 1.265 | 4.131 | 38.860 | 2.15e-05 | 0.012487411 |
| ENSDARG00000005085 | *ggctb* | 0.505 | 5.399 | 35.011 | 2.43e-05 | 0.013723445 |
| ENSDARG00000039754 | *xpc* | -0.624 | 4.628 | 34.711 | 2.55e-05 | 0.014010615 |

| | | | | | | |
|---|---|---|---|---|---|---|
| ENSDARG00000069951 | *eef1a1l2* | -0.609 | 5.125 | 33.215 | 3.24e-05 | 0.017088134 |
| ENSDARG00000069377 | *si:dkey-242g16.2* | -0.766 | 4.696 | 33.161 | 3.27e-05 | 0.017088134 |
| ENSDARG00000036045 | *penkb* | 1.025 | 2.579 | 32.876 | 3.43e-05 | 0.017282185 |
| ENSDARG00000095294 | *si:dkey-200l5.4* | -1.427 | 2.485 | 32.795 | 3.48e-05 | 0.017282185 |
| ENSDARG00000038465 | *stmn3* | -0.612 | 4.781 | 32.440 | 3.69e-05 | 0.017904601 |
| ENSDARG00000068951 | *si:ch211-219a15.4* | 2.843 | 0.624 | 33.394 | 3.81e-05 | 0.018078068 |
| ENSDARG00000053130 | *pcp4a* | -0.661 | 4.852 | 31.938 | 4.01e-05 | 0.018606096 |
| ENSDARG00000094990 | *si:dkey-91f15.1* | 1.648 | 3.381 | 35.342 | 4.25e-05 | 0.019295365 |
| ENSDARG00000038359 | *enosf1* | 0.560 | 4.965 | 30.600 | 5.04e-05 | 0.022385334 |
| ENSDARG00000036186 | *mbpa* | -0.513 | 6.301 | 30.383 | 5.23e-05 | 0.022761232 |
| ENSDARG00000007769 | *sult5a1* | 2.191 | 1.616 | 33.786 | 5.35e-05 | 0.022820274 |
| ENSDARG00000005372 | *camk4* | -0.575 | 4.518 | 34.802 | 5.63e-05 | 0.023516933 |
| ENSDARG00000017490 | *cel.1* | -1.327 | 9.128 | 42.716 | 6.18e-05 | 0.025296181 |
| ENSDARG00000088711 | *lgals1l1* | 1.331 | 4.044 | 36.251 | 6.49e-05 | 0.026064889 |
| ENSDARG00000023082 | *krt1-19d* | -0.520 | 7.295 | 28.834 | 6.88e-05 | 0.027120305 |
| ENSDARG00000034503 | *per2* | -0.502 | 7.783 | 28.599 | 7.18e-05 | 0.027774121 |
| ENSDARG00000062788 | *irg1l* | 1.759 | 2.747 | 33.320 | 7.73e-05 | 0.029316922 |
| ENSDARG00000100515 | *dusp1* | -0.735 | 7.465 | 31.726 | 7.86e-05 | 0.029316922 |
| ENSDARG00000077982 | *elf3* | 0.918 | 3.989 | 29.039 | 8.00e-05 | 0.029316922 |
| ENSDARG00000052779 | *zgc:153932* | 3.735 | 3.282 | 45.407 | 8.54e-05 | 0.029942295 |
| ENSDARG00000039393 | *si:ch211-240l19.5* | -0.628 | 5.495 | 27.647 | 8.55e-05 | 0.029942295 |
| ENSDARG00000023228 | *vsnl1a* | -0.439 | 6.721 | 39.331 | 8.60e-05 | 0.029942295 |
| ENSDARG00000018542 | *hapln4* | -1.405 | 1.140 | 27.273 | 9.17e-05 | 0.031397972 |
| ENSDARG00000018997 | *cplx2l* | -0.433 | 6.652 | 34.489 | 9.69e-05 | 0.032634538 |
| ENSDARG00000025301 | *gfap* | 0.491 | 7.693 | 26.790 | 1.00e-04 | 0.033296277 |
| ENSDARG00000090297 | *ldlrad2* | 0.977 | 2.169 | 26.435 | 1.07e-04 | 0.035067226 |
| ENSDARG00000058992 | *cers2b* | 0.574 | 4.176 | 29.458 | 1.10e-04 | 0.035237692 |
| ENSDARG00000038609 | *mpz* | -0.754 | 4.901 | 27.377 | 1.14e-04 | 0.036081227 |
| ENSDARG00000058206 | *si:ch211-153b23.5* | 0.554 | 5.018 | 25.854 | 1.20e-04 | 0.037009979 |
| ENSDARG00000002758 | *dedd1* | -0.465 | 7.515 | 25.840 | 1.21e-04 | 0.037009979 |
| ENSDARG00000026726 | *anxa1a* | -0.465 | 8.092 | 25.727 | 1.23e-04 | 0.037097551 |
| ENSDARG00000003146 | *si:dkey-27m7.4* | -0.914 | 3.146 | 25.582 | 1.27e-04 | 0.037097551 |
| ENSDARG00000058332 | *KRT18 (1 of many)* | -0.853 | 2.739 | 25.547 | 1.28e-04 | 0.037097551 |
| ENSDARG00000039522 | *tubb2* | -0.424 | 5.420 | 30.844 | 1.28e-04 | 0.037097551 |
| ENSDARG00000098256 | *si:ch73-329n5.6* | 3.071 | 0.161 | 25.221 | 1.36e-04 | 0.038906766 |
| ENSDARG00000045269 | *pts* | 0.649 | 3.477 | 25.212 | 1.43e-04 | 0.040450174 |
| ENSDARG00000004748 | *zgc:100868* | 0.981 | 6.686 | 32.601 | 1.48e-04 | 0.041174301 |
| ENSDARG00000016238 | *arl6ip5b* | -0.445 | 5.497 | 24.511 | 1.57e-04 | 0.043044942 |
| ENSDARG00000086705 | *prob1* | 0.901 | 2.738 | 24.384 | 1.61e-04 | 0.043586532 |
| ENSDARG00000055752 | *npas4a* | -1.327 | 6.570 | 34.241 | 1.65e-04 | 0.043786913 |
| ENSDARG00000030980 | *csrp1b* | -0.684 | 3.584 | 27.065 | 1.66e-04 | 0.043786913 |

| | | | | | | |
|---|---|---|---|---|---|---|
| ENSDARG00000043932 | *stmn4l* | 0.433 | 5.347 | 28.293 | 1.78e-04 | 0.045886342 |
| ENSDARG00000070442 | *si:ch211-113g11.6* | -0.424 | 6.260 | 33.027 | 1.78e-04 | 0.045886342 |
| ENSDARG00000039502 | *eef1a1a* | -0.430 | 6.668 | 37.536 | 1.84e-04 | 0.046899431 |

*Notes:* Log2 fold change reported (logFC) in which + indicates a gene that is up-regulated in DomA-exposed fish relative to controls, and - indicates a gene that is down-regulated in DomA-exposed fish relative to controls. False discovery rate (FDR) applied for multiple testing correction using the Benjamini-Hochberg method.

**Supplemental Table 27: Differentially expressed genes in DomA-exposed fish at 7 dpf (exposed at 2 dpf then tissue extracted at 7 dpf)**

| ENSEMBLgene | Gene name | Gene description | logFC | logCPM | F | PValue | FDR | DGE at 3 dpf? |
|---|---|---|---|---|---|---|---|---|
| ENSDARG00000043697 | *nefmb* | neurofilament, medium polypeptide b | -1.056 | 5.258 | 66.029 | 7.46E-07 | 0.0083 | Y |
| ENSDARG00000031588 | *si:dkey-239b22.1* | si:dkey-239b22.1 | 1.573 | 3.901 | 67.996 | 7.97E-07 | 0.0083 | Y |
| ENSDARG00000096849 | *si:dkey-16p21.8* | si:dkey-16p21.8 | -0.530 | 7.452 | 46.685 | 6.97E-06 | 0.0433 | N |
| ENSDARG00000009505 | *prelid3b* | PRELI domain containing 3 | -0.550 | 7.584 | 42.332 | 8.30E-06 | 0.0433 | N |
| ENSDARG00000038716 | *casq1a* | calsequestrin 1a | -0.681 | 7.079 | 40.746 | 1.09E-05 | 0.0454 | N |
| ENSDARG00000097959 | *si:dkey-248g15.3* | si:dkey-248g15.3 | -0.555 | 5.734 | 36.912 | 1.81E-05 | 0.0491 | N |
| ENSDARG00000012426 | *neflb* | neurofilament, light polypeptide b | -0.682 | 4.325 | 36.005 | 2.08E-05 | 0.0491 | Y |
| ENSDARG00000028507 | *itgb4* | integrin, beta 4 | -0.656 | 6.650 | 37.036 | 2.25E-05 | 0.0491 | N |
| ENSDARG00000057571 | *pgam2* | phosphoglycerate mutase 2 (muscle) | -0.478 | 6.553 | 37.467 | 2.25E-05 | 0.0491 | N |
| ENSDARG00000004187 | *zgc:122979* | zgc:122979 | -0.609 | 5.832 | 35.445 | 2.35E-05 | 0.0491 | Y |

*Notes:* + indicates a gene that is up-regulated in DomA exposed fish relative to controls, - indicates a gene that is down-regulated in DomA exposed fish relative to controls.
